# Structural basis for the ligand recognition and signaling of free fatty acid receptors

**DOI:** 10.1101/2023.08.20.553924

**Authors:** Xuan Zhang, Abdul-Akim Guseinov, Laura Jenkins, Kunpeng Li, Irina G. Tikhonova, Graeme Milligan, Cheng Zhang

## Abstract

Free fatty acid receptors 1-4 (FFA1-4) are class A G protein-coupled receptors (GPCRs). FFA1-3 share substantial sequence similarity whereas FFA4 is unrelated. Despite this FFA1 and FFA4 are activated by the same range of long chain fatty acids (LCFAs) whilst FFA2 and FFA3 are instead activated by short chain fatty acids (SCFAs) generated by the intestinal microbiota. Each of FFA1, 2 and 4 are promising targets for novel drug development in metabolic and inflammatory conditions. To gain insights into the basis of ligand interactions with, and molecular mechanisms underlying activation of, FFAs by LCFAs and SCFAs, we determined the active structures of FFA1 and FFA4 bound to the polyunsaturated LCFA docosahexaenoic acid (DHA), FFA4 bound to the synthetic agonist TUG-891, as well as SCFA butyrate-bound FFA2, each complexed with an engineered heterotrimeric G_q_ protein (miniG_q_), by cryo-electron microscopy. Together with computational simulations and mutagenesis studies, we elucidated the similarities and differences in the binding modes of fatty acid ligands with varying chain lengths to their respective GPCRs. Our findings unveil distinct mechanisms of receptor activation and G protein coupling. We anticipate that these outcomes will facilitate structure-based drug development and underpin future research to understand allosteric modulation and biased signaling of this group of GPCRs.

## Introduction

Free fatty acids are bioactive lipids comprising a carboxylic acid head group and an aliphatic hydrocarbon chain with various lengths. In humans, and many other species, they can activate a group of G protein-coupled receptors (GPCRs) including free fatty acid receptors 1-4 (FFA1-4 receptors) and GPR84 to regulate metabolic homeostasis and immunity ^1–3^. Among them, FFA1 (GPR40) and FFA4 (GPR120) mainly sense long-chain fatty acids (LCFAs) with more than 12 carbons while FFA2 (GPR43) and FFA3 (GPR41) primarily sense short-chain fatty acids (SCFAs) with less than 6 carbons ^3,4^. Representative LCFA ligands of FFA1 and FFA4 include ω-3 and ω-6 polyunsaturated fatty acids (PUFAs) ^5,6^. Meanwhile, SCFA ligands of FFA2 and FFA3 are mainly produced in the gut as products of microbiota-mediated fermentation and include acetate, propionate, and butyrate ^7–12^.

FFAs play critical roles in both immunity and metabolism ^3,4,10^. FFA1 signaling induced by LCFAs in pancreatic β cells can facilitate insulin secretion ^13^, making it a promising drug target for type 2 diabetes mellitus (T2D) ^14,15^. Although the FFA1-selective agonist TAK-875, also named fasiglifam, exhibited promising antidiabetic effects in clinical studies, it failed in phase III trials due to liver toxicity ^16^. However, other FFA1 activators are still being pursued for the treatment of T2D ^14^. FFA4, which has been described as the ω-3 PUFA receptor, is highly expressed in adipose tissue and macrophages. It mediates anti-inflammatory and other beneficial effects of ω-3 PUFAs such as docosahexaenoic acid (DHA) in those tissues and cells ^3–5,17–21^. FFA4 is also considered as a new drug target for diabetes ^21–23^. FFA4 selective or FFA1/FFA4 dual agonists ^24–26^ may hold the promise of becoming a new class of antidiabetic drugs with additional anti-inflammatory benefits ^22,26^. Interestingly, in addition to their functions in metabolism and immunity, both FFA1 and FFA4, particularly FFA4, have been suggested to function as lipid taste receptors ^27–29^. On the other hand, FFA2 and FFA3 are expressed in adipocytes and a range of immune cells. Their unique ligand preference of SCFAs produced by the fermentation of dietary fiber in the lower gut has led to intensive research on their roles at the interface of host and gut microbiota ^30,31^. Previous studies suggested that many beneficial effects of gut microbiota on the host, including the resolution of inflammation ^32^, suppression of fat accumulation ^33^, and protection from viral and bacterial pathogens ^34,35^, are mainly mediated by the SCFA-FFA2 signaling axis. Therefore, FFA2 and FFA3, especially FFA2, are considered as new promising therapeutic targets for metabolic disorders, including obesity and diabetes, and inflammatory diseases ^3,10,36–38^. However, compared to FFA1 and FFA4, fewer synthetic ligands have been reported for FFA2 and FFA3, which may suggest certain obstacles in developing small molecule ligands for these two SCFA receptors.

Phylogenetic analysis suggests that FFA4 diverged early from other FFAs (**Fig. S1**). As a result, while FFA1-3 are structurally related with high sequence similarity, FFA4 shares very little sequence similarity with FFA1-3 ^39^. This implies distinct ligand recognition and signaling mechanisms for FFA4 and other FFAs. Regarding G protein coupling, FFA1 is a highly promiscuous GPCR that is capable of coupling to all four G protein families: G_s_, G_i/o_, G_q/11_, and G_12/13_ ^39,40^. FFA2 and FFA4 both can signal through G_i/o_ and G_q/11_ ^10,39–41^. For FFA4, a human splice variant (FFA4^Long^) has been identified with an additional 16-amino acid segment in the intracellular loop 3 (ICL3). This isoform is unable to induce G_q/11_ signaling but is capable of coupling to β-arrestins ^42,43^. Crystal structures of highly engineered forms of FFA1 bound to synthetic agonists including TAK-875 and positive allosteric modulators (PAMs) have been reported ^44–46^, and the receptor in those structures stayed in the inactive state. To understand the molecular mechanisms by which LCFAs and SCFAs act on and activate their FFAs, we determined active structures of DHA-bound FFA1, butyrate-bound FFA2, and DHA-bound FFA4 in complex with an engineered heterotrimeric G_q_ protein (miniG_q_) ^47^ by cryo-electron microscopy (cryo-EM). To further examine how FFA4 recognizes different ligands, we also solved a cryo-EM structure of the FFA4-miniGq complex with the most widely employed synthetic FFA4 agonist, TUG-891 ^48^. These structures revealed diverse modes of ligand recognition by FFAs. Together with computational simulations and mutagenesis studies, these studies highlight similarities and differences in modes of binding of the fatty acid ligands of varying chain length to their corresponding GPCRs.

## Results

### Overall structures of FFA signaling complexes

We used the wild type human FFA1, FFA2, and FFA4 in our structural studies. For FFA4, there are two isoforms and we chose the canonical short form since FFA4^Long^ doesn’t couple to G_q/11_ ^43^. The miniG_q_ protein contains an engineered miniG_αq_ subunit ^49^ with the N-terminal 35 amino acids replaced by their corresponding residues in G_αi_. The same miniG_q_ protein has been successfully used to obtain cryo-EM structures of several other G_q_-coupled GPCRs ^47,50–53^. To further stabilize the FFA-miniG_q_ complexes, we assembled the complexes using the NanoBit tethering strategy in insect Sf9 cells ^54^ together with an antibody fragment, scFv16, which has been developed previously to stabilize the G_i_ heterotrimer ^55^. The structures of DHA-bound FFA1 and FFA4, TUG-891-bound FFA4, and butyrate-bound FFA2 with miniG_q_ were determined to overall resolutions of 3.4 Å, 3.2 Å, 3.1 Å, and 3.1 Å, respectively (**Fig. 1a, Fig. S2-5, Table S1-2**).

**Figure 1.**
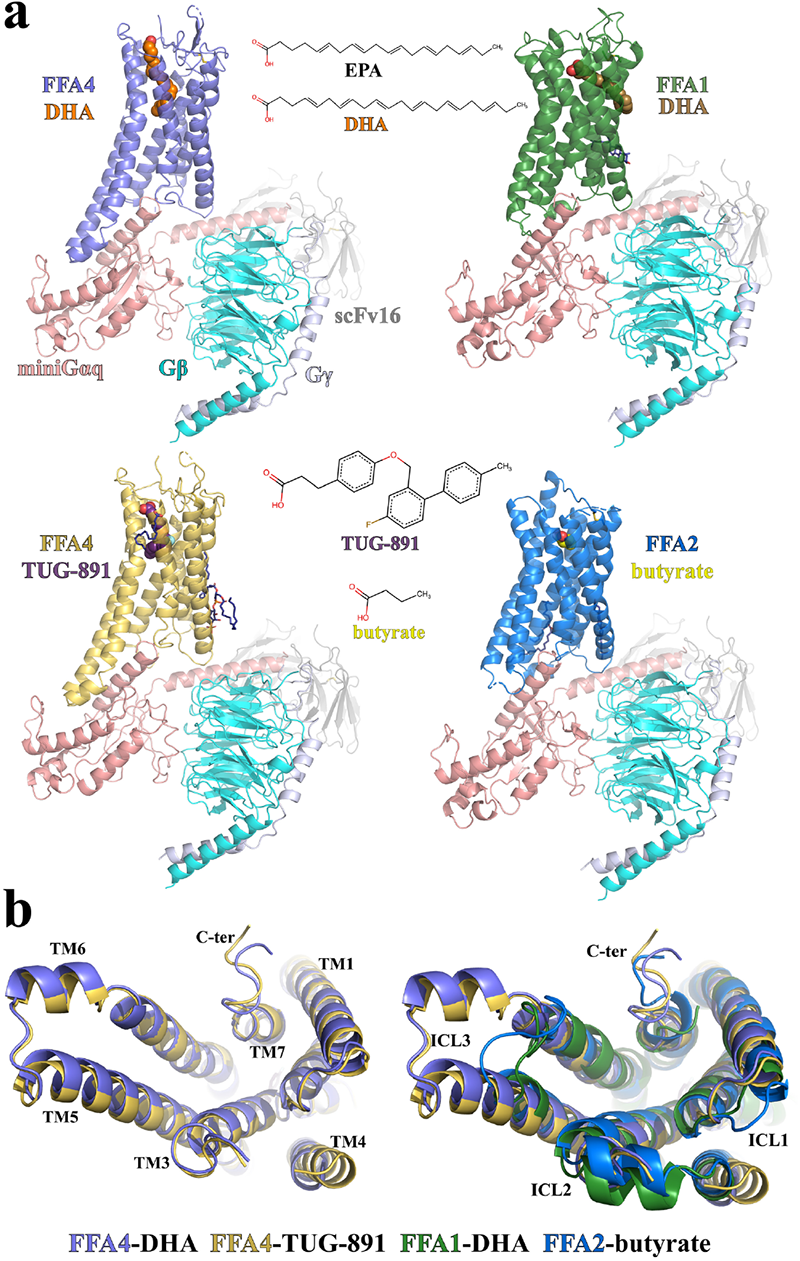
Overall structures of FFA1, FFA2, and FFA4 signaling complexes. **(a)** Overall structures of DHA-FFA4 (slate), DHA-FFA1 (green), TUG-891-FFA4 (dark yellow) and butyrate-FFA2 (blue), each in complex with miniG_q_, are shown, as are the chemical structures of the bound ligands. miniG_αq_, G_β_ and G_γ_ subunits are colored in salmon, cyan and light blue, respectively. ScFv16 is colored grey. The LCFA eicosapentaenoic acid (EPA) is also shown for comparison to DHA (see main text for discussion). **(b)** Comparison of the above structures as seen from the intracellular face.

The majority of residues in the three FFAs, miniG_q_, and scFv16 were modeled based on the robust cryo-EM density maps. We also modeled several cholesterol and lipid molecules to fit strong density maps surrounding the transmembrane domains (TMDs) of FFA2 and FFA4 (**Fig. S6**). In the two structures of FFA4, the density of the intracellular region of transmembrane helix 4 (TM4) and intracellular loop 2 (ICL2) is relatively weak, indicating a high degree of flexibility (**Fig. 1b**). In contrast, ICL2 forms a helical structure in both FFA1 and FFA2 (**Fig. 1b**). A large part of the extracellular loop 2 (ECL2) of FFA2 is also not modeled due to weak density. Noticeably, in the structures of all three receptors, no helix 8 after TM7 was modeled due to very weak density. This suggests that the C-terminal region after TM7 in all three receptors is highly mobile when coupled with G proteins.

As for the ligands, the density maps for TUG-891 and DHA in FFA4 were sufficiently clear to enable modeling of the entire ligands (**Fig. S2-3**). Additionally, the density of butyrate in FFA2 was also strong (**Fig. S5**). However, due to the small size of butyrate and the limited resolution of the cryo-EM map, functional data was necessary to complement cryo-EM map information for accurate ligand modeling. In the case of FFA1, we modeled DHA in the orthosteric site based on a partial density map (**Fig. S4**). Further discussion on this topic will be provided in the subsequent content.

### Binding of DHA and the synthetic agonist TUG-891 to FFA4

Both DHA and TUG-891 bind to a pocket formed among the extracellular regions of TM3-7 of FFA4 (**Fig. 2a**). The amino-terminal (N-terminal) region of FFA4 preceding TM1 folds inside the TMD and directly interacts with the ligands (**Fig. 2a**), resulting in almost complete shielding of the ligand-binding pocket from the extracellular milieu (**Fig. 2b**). This is similar to the N-terminal region of DP2, a GPCR that binds the fatty acid ligand prostaglandin D_2_ (PGD_2_), and that forms a well-folded structure that participates in ligand interactions ^56,57^.

**Figure 2.**
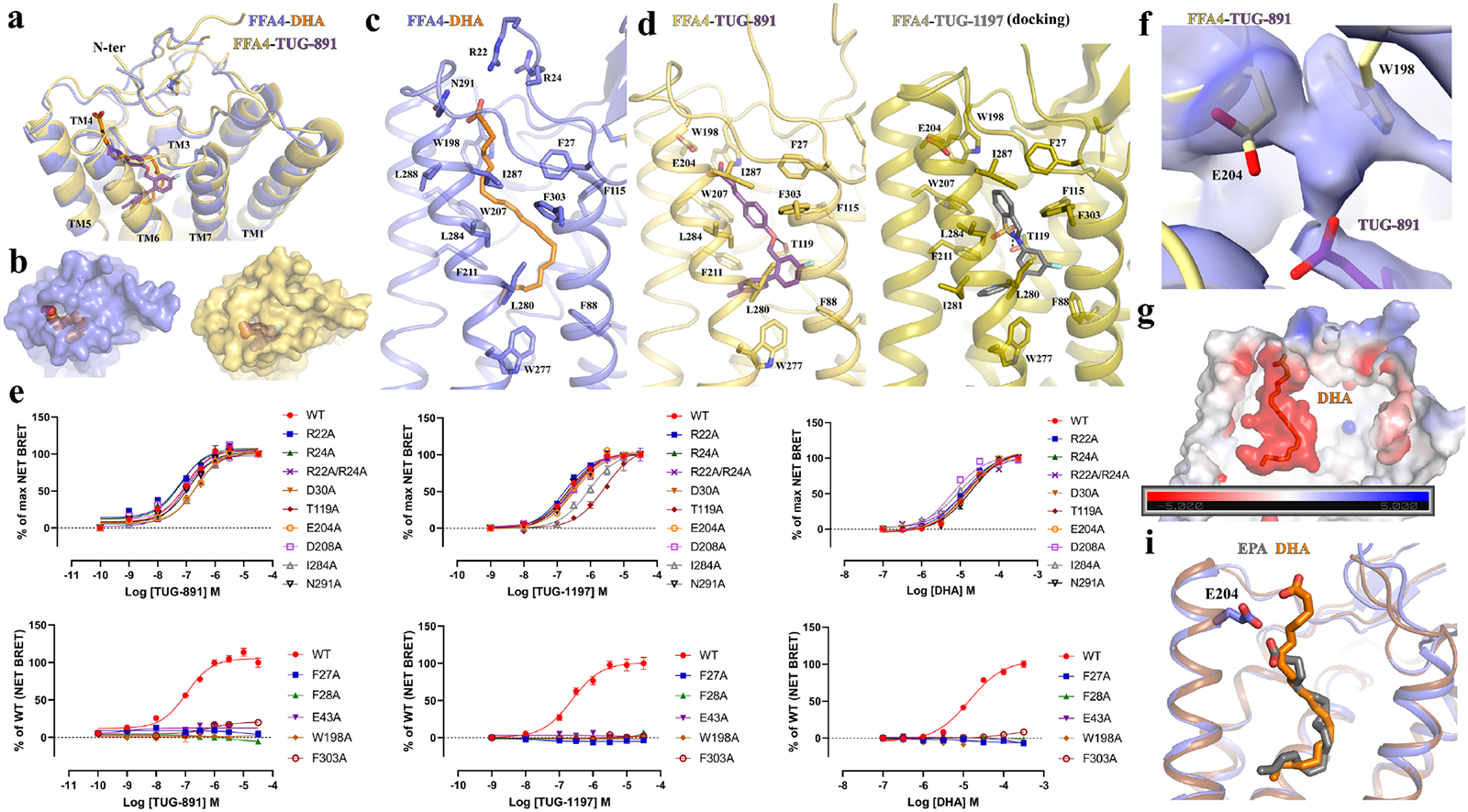
Ligand binding in FFA4. **(a-d)** Details of the interactions of DHA (orange) and TUG-891 (purple) with FFA4. Panel **a** illustrates the general positions of the two ligands whilst **b** highlights the closed nature of the occupied ligand binding pockets. Details of key residues of the binding pockets are highlighted for DHA (**c**) and TUG-891 (**d, left**). TUG-1197 docked into the FFA4 structure (**d, right**) highlights the important role of T119 and the similarity of the binding mode of TUG-1197 and TUG-891 at the bottom of the pocket. **(e)** Various point mutants of FFA4 generated and assessed for the ability of each of TUG-891, TUG-1197 and DHA to promote interactions with arrestin-3. See **Fig. S7** for quantitation. In concert with the large-scale mutagenesis studies reported previously ^59^, this provides a comprehensive analysis of the orthosteric binding pocket of FFA4. (**f**) Continuous electron density observed between W198 and E204. The cryo-EM map is contoured at the level of 0.13. (**g**) Negative charge potential of the FFA4 binding pocket with DHA. **(h)** Additional length of DHA compared to EPA and the position of the DHA carboxylate above and beyond E204.

The cryo-EM density is strong for the ω-3 unsaturated chain of DHA, while the density for the carboxylate group is comparatively weaker. Nevertheless, we were able to model the entire DHA molecule that well fits the density. Our structure revealed that the ω-3 unsaturated chain of DHA adopts a ‘L’ shape binding pose, which enables it to penetrate deeply into a binding pocket that is rich in aromatic residues (**Fig. 2a**). The carboxylate head group of DHA, on the other hand, extends outward towards the extracellular milieu (**Fig. 2a**). The six carbon-carbon double bonds in DHA are surrounded by, and potentially form, extensive π-π interactions with aromatic residues F27^N^, F28^N^, F88^2.53^, F115^3.29^, W198^ECL2^, W207^5.38^, F211^5.42^, W277^6.48^, and F303^7.35^ (superscripts represent Ballesteros-Weinstein numbering ^58^) (**Fig. 2c**). Hydrophobic residues I280^6.51^, I284^6.55^, I287^6.58^, and L288^6.59^ lining one side of TM6, together with M118^3.32^, L196^ECL2^, and I287^6.58^ form additional hydrophobic interactions with DHA to further stabilize lipid binding (**Fig. 2c**).

TUG-891 (3-(4-((4-fluoro-4-methyl-[1,1-biphenyl]-2-yl)methoxy)phenyl)-propanoic acid) binds to FFA4 in a similar ‘L’ shape binding pose as DHA, overlapping extensively with the ω-3 chain of DHA and with the ortho-biphenyl moiety of TUG-891 defining the bottom of the binding pocket (**Fig. 2a**). This leads to the observation of similar sets of hydrophobic and π-π interactions between the three benzene rings of TUG-891 and FFA4 (**Fig. 2d**). Mutagenesis studies we performed previously are in accord with these observations. Alteration of F88^2.53^, F115^3.29^, W207^5.38^, F211^5.42^, W277^6.48^ (each to A) and F303^7.35^ (to H) (and also here to A, **Fig. 2e**) resulted in either a complete lack of response to TUG-891 or a greater than 100-fold reduction in potency ^59^. The mutation W198A also lacked response to both TUG-891 and DHA (**Fig. 2e, Fig. S7**). However, although mutations to Ala of F27^N^ and F28^N^ each lacked response to TUG-891 (**Fig. 2e, Fig. S7**), these results could not be interpreted because, although well expressed following transient transfection in HEK293 cells (**Fig. S7c**), each of these mutants failed to reach the cell surface (**Fig. S7d**). Mutation to Ala of either I280^6.51^ or I284^6.55^ also produced a greater than 100-fold (I280A) ^59^ or a more modest but still significant (I284A) reduction in potency for TUG-891 (**Fig. 2e, Fig. S7a**). We previously observed a similar pattern of effects of these mutations for the ω-3 PUFA α-linolenic acid (*all*-*cis*-9,12,15-octadecatrienoic acid) ^59^. Although not altering the potency of DHA (**Fig. 2e, Fig. S7a**), a notable feature of the I284A mutant was that it reduced the efficacy of DHA, such that it acted as a partial agonist compared to TUG-891 at this mutant **(Fig. S7b**), a feature that was not observed for the other mutants studied.

An additional difference in the binding mode of TUG-891 and DHA is that the linking ether oxygen of TUG-891 forms a hydrogen bond with T119^3.33^, which is absent in the DHA-bound FFA4 (**Fig. 2c and d**). Notably, mutation of T119A significantly impaired the ability of TUG-891 to activate FFA4, in both β-arrestin interaction **(Fig. S7a**) and, particularly, G_q_-mediated Ca^2+^ elevation assays ^59^, indicating an important role of this hydrogen bond in TUG-891 binding and function. No such effects of this mutation on the function of either DHA (**Fig. 2e, Fig. S7a**) or α-linolenic acid ^59^ was observed. A distinct group of sulphonamide-based FFA4 agonists have been reported ^39,60^. Among them, TUG-1197 (2-(3-(pyridin-2-yloxy)phenyl)-2,3-dihydrobenzo[d]isothiazole 1,1-dioxide)) showed a large, greater than 10-fold, reduction in potency at the T119A mutant and greatly reduced efficacy in comparison to TUG-891 (**Fig. 2e, Fig. S7a-b**). Docking of this ligand to the obtained structures of FFA4 suggests a similar binding pose as TUG-891 and a clear interaction of the sulphonamide, which overlaps location with the ether oxygen of TUG-891, with T119 (**Fig. 2d**).

Intriguingly, in the structure, the carboxylate group of the phenyl-propanoic acid of TUG-891 is positioned in proximity to E204^5.35^ and W198^ECL2^ (**Fig. 2d**). Remarkably, the cryo-EM density between E204^5.35^ and W198^ECL2^ appears to be continuous (**Fig. 2f**), which raises the possibility that a water molecule may be present between these two residues to facilitate extensive polar interactions between the carboxylate group of TUG-891 and E204^5.35^ and W198^ECL2^ of FFA4. The mutant E204A modestly but significantly reduced potency of TUG-891 but not of DHA (**Fig. S7a**) whereas a W198A mutant was not activated by either TUG-891 or DHA (**Fig. 2e, Fig. S7a).**

The carboxylate group of DHA, which is associated with weak cryo-EM density, is modeled close to polar residues R22^N^ and R24^N^ from the N-terminal region and N291^ECL3^ from ECL3 (**Fig. 2c**). However, single mutations R22A and R24A or the double mutation R22A/R24A did not reduce the potency of DHA or either of the synthetic agonists TUG-891 and TUG-1197 (**Fig. 2e**, **Fig. S7a**). Therefore, the binding of DHA to FFA4 is mainly driven by hydrophobic and π-π interactions. Nevertheless, it is to be noted that the overall binding pocket of DHA exhibits a negatively charged potential, which may help to position the carboxylate group of DHA at the extracellular surface (**Fig. 2g**). A similar charge interaction-facilitated lipid recognition mechanism has also been suggested for other lipid GPCRs including prostaglandin E_2_ (PGE_2_) receptors and lysophospholipid GPCRs ^56,57^. However, although DHA adopts a similar binding orientation as PGE_2_ and lysophospholipids, their binding sites are located differently (**Fig. S8a**). The pockets of PGE_2_ and lysophospholipids form among TM1-TM2-TM3-TM7 or TM2-TM3-TM5-TM6-TM7, while the pocket of DHA in FFA4 forms among TM3-TM4-TM5-TM6-TM7 (**Fig. S8a**). To the best of our knowledge, no other lipid GPCRs have been shown to have lipid binding pockets at similar locations as that of DHA in FFA4.

During the preparation of our manuscript, other research groups published structures of FFA4 bound to several LCFAs, including eicosapentaenoic acid (EPA), an ω-3 PUFA, and TUG-891 ^61^. The structure of EPA bound to FFA4 showed a highly similar binding pose to that of DHA observed in our structure, especially with regard to their ω-3 chains (**Fig. 2h**). However, DHA is two carbons longer than EPA and contains an additional double bond. As a consequence, the carboxylate group of DHA extends further towards the extracellular surface above E204^5.35^, while the carboxylate group of EPA is located below E204^5.35^ (**Fig. 2h**). Furthermore, we observed a slightly different binding mode of TUG-891 in our cryo-EM structure, which is strongly supported by clear cryo-EM density, compared to the published structure (**Fig. S8b**). The overall position of TUG-891 in the published structure is closer to the extracellular surface compared to that in our structure (**Fig. S8b**). As a result, the carboxylate group of TUG-891 in our structure is too distant from N291^ECL3^ to form a hydrogen bond. Consistent with this, we did not observe an effect of the N291A mutant on the potency of TUG-891 (**Fig. 2e, Fig S7a**) and such a mutant was not reported in the published study ^61^. In addition, the hydrogen bond between the ether oxygen of TUG-891 and T119^3.33^ in our structure is absent in the published structure (**Fig. S8b**). The discrepancies in TUG-891 binding in the two structures may indicate a high degree of mobility of TUG-891 in FFA4. Nevertheless, despite these discrepancies, our mutagenesis studies demonstrated the important role of T119 in the action of TUG-891 (**Figs. S7a, S8b**) and the positioning of the ortho-biphenyl, which is key feature of TUG-891 and related synthetic FFA4 agonists, is entirely in accord with our earlier mutagenesis studies ^59^.

### Distinct mechanisms of DHA recognition by FFA1 and FFA4

Despite their similar ligand recognition profiles as LCFA receptors, FFA1 and FFA4 exhibit little sequence similarity and a distant phylogenetic relationship (**Fig. S1**). To investigate whether FFA1 utilizes a distinct mechanism to recognize DHA, we sought to obtain a cryo-EM structure of miniGq-coupled FFA1 bound to DHA. However, while the overall resolution of the structure reached 3.4 Å (**Fig. S4**), modeling DHA proved to be challenging.

Previous structural studies on FFA1 with synthetic agonists and allosteric modulators ^44–46^ have identified two binding sites: ‘Site 1’ is located in the extracellular region within the 7TM as the putative orthosteric site for synthetic agonists TAK-875 ^46^ and MK-8666 ^44^, while ‘Site 2’ is located on the surface of the 7TM above ICL2 as the allosteric site for the synthetic ago-positive allosteric modulators (ago-PAMs) AP8 ^44^ and Compound 1 (3-benzyl-4-(cyclopropyl-(4-(2,5-dichlorophenyl)thiazol-2-yl)amino)-4-oxobutanoic acid) ^45^. Similar allosteric sites have also been identified for C5a receptor ^62^ and β2-adrenergic receptor ^63^. Evidence from a previous study showed that TAK-875 exhibited positive cooperativity with the LCFA ligand of FFA1 γ-linolenic acid (γ-LA), suggesting that ‘Site 1’ is not the primary site for γ-LA ^64^.

In our structure, we observed weak density in ‘Site 1’ for DHA (**Fig. 3a).** We further performed local refinement focusing on the receptor to improve this density, which allowed us to model the part of DHA from C1 to C8 together with the carboxylate group in ‘Site 1’ (**Fig. 3a**). The region of DHA from C9 to C22 was assigned with zero occupancy in the structure since it couldn’t be well modeled. In our structural model, the carboxylate group of DHA forms salt bridges with R183^5.39^ and R258^7.35^, while the carbon chain from C1 to C8 forms hydrophobic and π-π interactions with surrounding residues F87^3.33^, L138^4.57^, F142^4.61^, W174^ECL2^, and L186^5.42^ of FFA1 (**Fig. 3a**). A very recent study also reported a similar partial cryo-EM density for DHA in ‘Site 1’ of FFA1 ^65^. It is to be noted that the density of most residues in ‘Site 1’ after local refinement was sufficiently clear to allow unambiguous modeling, suggesting that the relatively weaker density of DHA was likely due to the high flexibility of DHA in this site.

**Figure 3.**
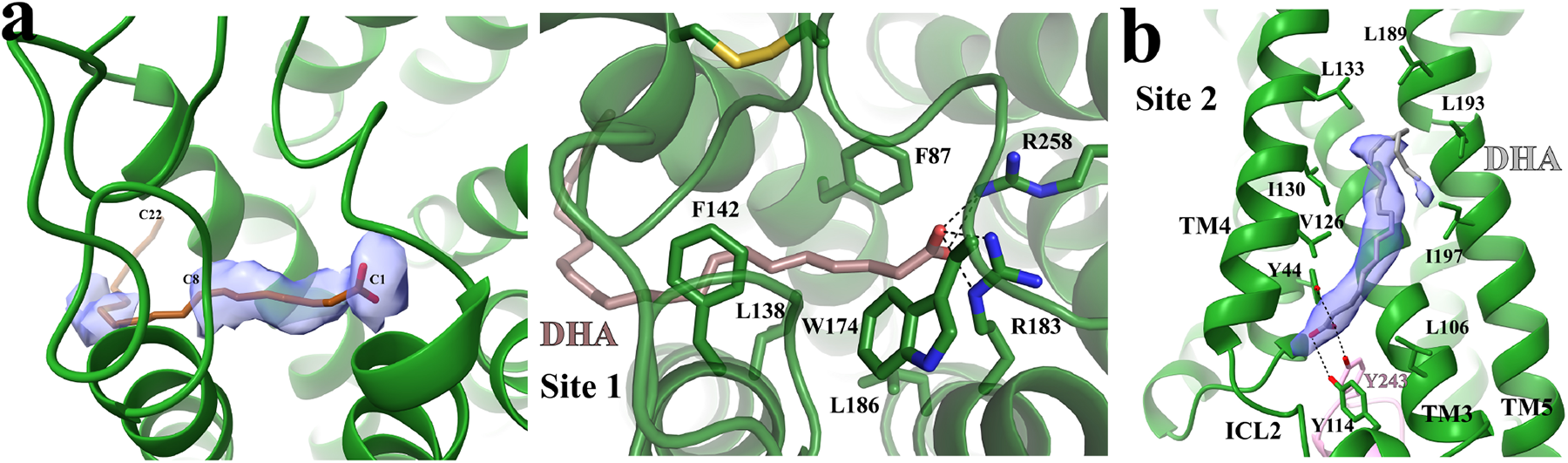
Potential DHA binding sites in FFA1. **(a)** DHA binding in ‘Site 1’. The partial cryo-EM density map of DHA colored in light blue in the left panel is contoured at the level of 0.07. C1, C8 and C22 atoms of DHA are labeled. The occupancy of DHA C9-C22 was assigned as zero due to a lack of density. The details of interactions between DHA and FFA1 in ‘Site 1’ are shown in the right panel. DHA is colored brown. **(b)** Putative DHA binding in ‘Site 2’. The strong cryo-EM density map in this site is contoured at the level of 0.12. The modeled DHA molecule is colored grey. Polar interactions are shown as black dashed lines. FFA1 is colored green.

Interestingly, strong density in ‘Site 2’ indicated the presence of a lipid molecule (**Fig. 3b**). It is possible that ‘Site 2’ is another binding site for DHA in FFA1. However, this site can also accommodate other lipid molecules, making it challenging to confirm it as the specific binding site for DHA. Nonetheless, since the cryo-EM sample contained a high concentration of DHA, it was the most prevalent lipid present. Thus, we fitted the density observed with a DHA molecule (**Fig. 3b**), and the binding pose of DHA in this site highly resembles that of the ago-PAM AP8 ^44,45^. The carboxylate group of DHA forms hydrogen bonds with two tyrosine residues Y44^2.42^ and Y114^ICL2^, while the carbon chain forms hydrophobic interactions with hydrophobic residues from TM3-5 of FFA1 (**Fig. 3b**). Interestingly, we also observed a hydrogen bond between DHA and Y243 from the α5 of mini-G_αq_, which is the major receptor interaction site in mini-G_αq_ (**Fig. 3b**).

In the bile acid receptor GPBAR, the endogenous lipid ligand bile acid binds to a site formed between TM3 and TM4 above ICL2, similar to ‘Site 2’ in FFA1 ^66^. However, bile acid can also bind to a more conventional orthosteric site located in the extracellular region of GPBAR ^66^. It is possible that DHA binds to FFA1 similarly to bile acid in GPBAR. ‘Site 1’ in FFA1 serves as the primary site for DHA, where DHA exhibits a high flexibility, while ‘Site 2’ in FFA1 serves as the secondary site for DHA.

Neither ‘Site 1’ nor ‘Site 2’ in FFA1 is conserved in FFA4, providing further evidence of the distant phylogenetic relationship between FFA1 and FFA4, despite their similar ligand preferences. In FFA4, the carboxylate group of DHA is positioned near the ligand entrance at the extracellular surface (**Fig. 2c**). In our previous studies on DP2, we observed that the prostaglandin PGD_2_ adopts a ‘polar-group-in’ binding pose in DP2 with its carboxylate group buried deep within the binding pocket, while another prostaglandin, PGE_2_, adopts a ‘polar-group-out’ binding pose in PGE_2_ receptors (EPs) with its carboxylate group positioned near the extracellular surface, and these two different binding poses of prostaglandins are facilitated by the distinct charge potentials of the binding pockets in DP2 and EPs ^56,57^. Similarly, DHA in FFA4 adopts a ‘polar-group-out’ binding pose in a negatively charged environment (**Fig. 2g**), although the role of the charge potential in DHA binding is not clear.

### Recognition of SCFAs by FFA2

As anticipated from the relatedness of FFA2 to FFA1 ^67^, the overall structures of these two receptors are similar. Also, butyrate and the carboxylate head group of DHA in ‘Site 1’ of FFA1 are very close if the structures of FFA1 and FFA2 are aligned (**Fig. 4a**). In the structure of FFA2-butyrate, the carboxylate group of butyrate is coordinated by a pair of adjacent arginine residues, R180^5.39^ and R255^7.35^ (**Fig. 4b**). These two arginine residues are highly conserved among FFA1-3 ^67,68^. Indeed, the same pair of residues in FFA1 interact with the carboxylate group of TAK-875 ^46^. In FFA2, mutating R180^5.39^ to other amino acids including Ala and Lys, as well as mutating R255^7.35^ to Ala, eliminates the response to SCFAs ^68^. Furthermore, the mutation to Ala of H242^6.55^, which interacts with R255^7.35^ to organize the binding pocket for the carboxylate of SCFAs (**Fig. 4b**), also abolishes SCFA function ^68^. These mutations eliminate the binding of SCFAs rather than simply affecting ligand function, as evidenced by the fact that SCFAs are unable to compete for binding with an FFA2 orthosteric antagonist, the affinity of which is only slightly reduced compared to the wild type receptor at each of the R180A, R255A, and H242A mutants of FFA2 ^69^. Interestingly, while R180^5.39^ and R255^7.35^ are conserved in FFA1, H242^6.55^ in FFA2 is replaced by N244^6.55^ in FFA1, which does not interact with R^5.39^ and R^7.35^.

**Figure 4.**
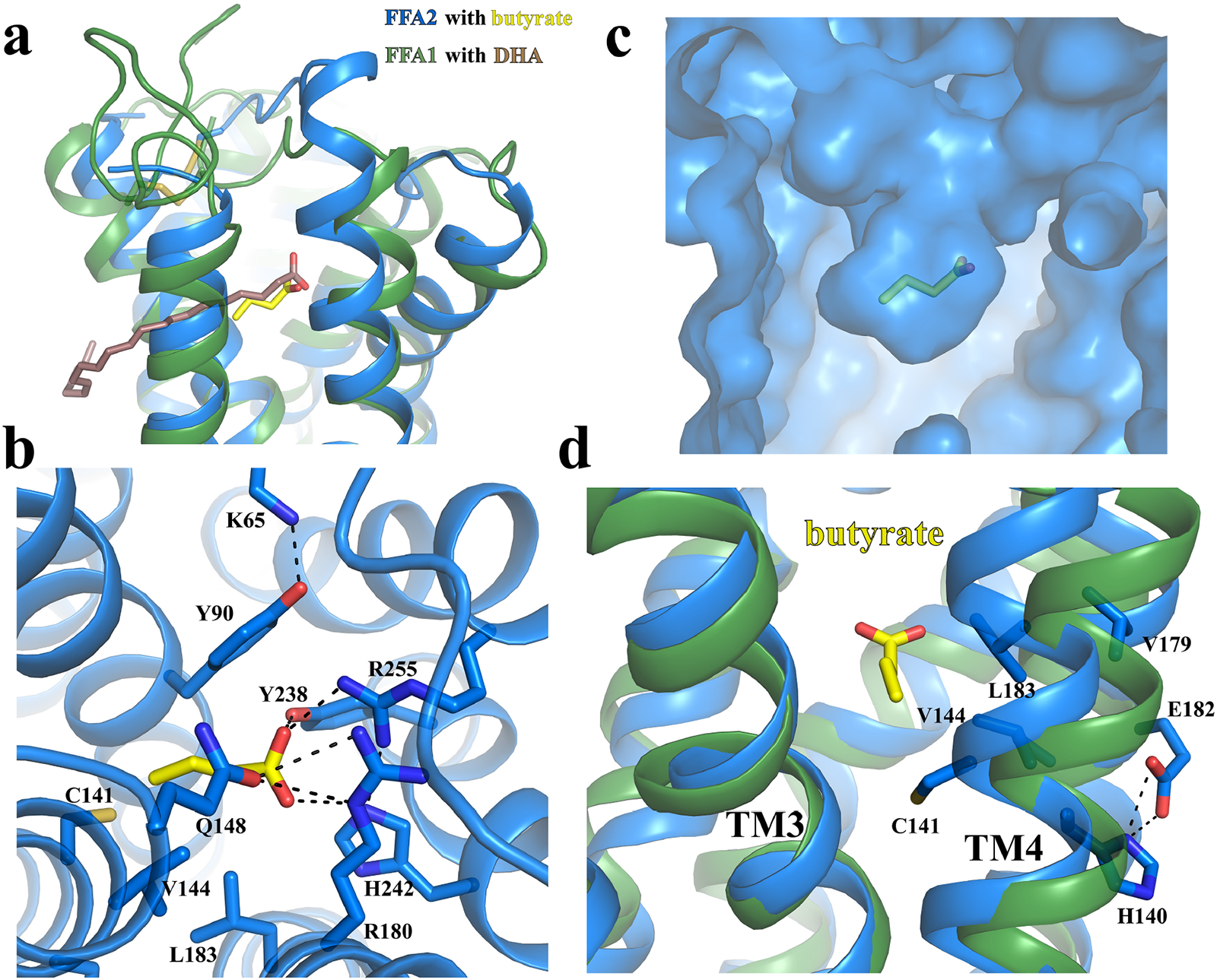
Recognition of butyrate by FFA2. **(a)** Structural alignment of FFA2-butyrate and FFA1-DHA. The carboxylate of butyrate occupies the equivalent position to the carboxylate of DHA. **(b)** Details of interactions between butyrate and FFA2. **(c)** Overall shape of the butyrate binding pocket. **(d)** Differences in location of TM3 and TM4 in FFA1 and FFA2. In all panels, FFA1 and FFA2 are colored in green and blue, respectively, whilst DHA and butyrate are colored in brown and yellow, respectively. Polar interactions are shown as black dashed lines.

In addition to R180^5.39^ and R255^7.35^, two other polar residues from ECL2, Q148^ECL2^ and Y165 ^ECL2^, are also close to butyrate (**Fig. 4b**). Y165 ^ECL2^ forms a hydrogen bond with the carboxylate of butyrate, while Q148^ECL2^ interacts with and potentially stabilizes R180^5.39^ (**Fig. 4b**). We have previously demonstrated that substituting Q148^ECL2^ to a glutamate residue results in a reduction of potency for SCFAs ^70^. It has an even more dramatic effect on larger synthetic FFA2 agonists including Compound 1 (3-benzyl-4-(cyclopropyl-(4-(2,5-dichlorophenyl)thiazol-2-yl)amino)-4-oxobutanoic acid), where the agonist function is all but ablated ^70^. This may be due to the repulsion of negatively charged carboxylate groups in the mutated E148 residue and agonists. Also, the mutation of Y165A resulted in a nearly 15-fold reduction of the potency of SCFAs ^70^, indicating the important role of this residue in ligand binding as well.

The opening between TM3 and TM4 of FFA1 that allows for binding of TAK-875 is closed in FFA2, resulting in a small pocket in FFA2 with limited space to accommodate the carbon chains of fatty acids (**Fig. 4c**). This may explain the selectivity of FFA2 for SCFAs over LCFAs. When comparing the structures of FFA1 and FFA2 coupled with miniGq, it becomes apparent that while TM3 aligns well between the two, TM4 in FFA2 shifts towards TM3 in comparison to FFA1 (**Fig. 4d**). H140^4.56^ in TM4 of FFA2 appears to play an important role in determining the selectivity of chain length of fatty acids. Whilst alteration of this residue to Ala reduces the potency of SCFAs, it enables binding and function of the C6 fatty acid caproate and, to a more modest degree, C8 caprylate ^68^. H140^4.56^ forms a hydrogen bond with E182^5.41^ in FFA2 (**Fig. 4d**), which is replaced by S185^5.41^ in FFA1. The longer side chain of E182^5.41^ may force H140^4.56^ together with TM4 to be positioned towards TM3. The H140^4.56^-E182^5.41^ pair also defines the size of the hydrophobic pocket involving C141^4.57^, V144^4.60^, V179^5.38^, and L183^5.42^ in FFA2 that accommodates the hydrophobic tail of butyrate (**Fig. 4d**). Notably, the mutations V179A and L183A do not change the potency of SCFAs but cause ∼10-fold reduction of the potency for larger synthetic FFA2 agonists Compound 1 and Compound 2 ((*R*)-3-(cyclopentylmethyl)-4-(cyclopropyl-(4-(2,6-dichlorophenyl)thiazol-2-yl)amino)-4-oxobutanoic acid) ^70^. C141^4.57^ is at the bottom of the SCFA binding pocket and forms van der Waals interactions with the last carbon of butyrate. This residue in bovine FFA2 is replaced by Gly. The C141G mutation in human FFA2 leads to an altered ligand preference for longer saturated and unsaturated C5 and C6 fatty acids, similar to that of bovine FFA2 ^71^.

In previous studies, two tyrosine residues in FFA2, Y90^3.29^ and Y238^6.51^, were suggested to participate in the binding of SCFAs ^69,70^. Mutating each residue to alanine significantly reduced the potency of SCFAs ^69,70^. In the structure of FFA2-butyrate, Y90^3.29^ forms direct hydrophobic interactions with the short chain of butyrate, while Y238^6.51^ forms hydrogen-bonding and cation-π interactions with the critical arginine residue R255^7.35^ (**Fig. 4b**), potentially stabilizing it in the appropriate position to interact with butyrate. Additionally, Y90^3.29^ is stabilized by a hydrogen bond with K65^2.60^ (**Fig. 4b**), and mutations of K65^2.60^ to either alanine or glutamate substantially reduce the potency of SCFAs ^72^. This indicates the important role of the K65^2.60^-Y90^3.29^ pair in SCFA binding. Interestingly, K65^2.60^ of human FFA2 is replaced by R65^2.60^ in mouse FFA2. It is notable that SCFAs including butyrate display lower potency at mouse FFA2 compared to human FFA2 ^73^. Consistent with this, a K65R mutation in human FFA2 resulted in a small but still significant reduction of potency of SCFAs ^72^.

### Molecular determinants of ligand recognition by FFAs revealed by MD simulations

To evaluate the stability and dynamics of the receptor-ligand interactions in FFA2 and FFA4, we performed molecular dynamics (MD) simulations of the two receptors in agonist-bound and unbound forms in a water-lipid bilayer. Guided by the protonation state prediction, D208^5.39^ was protonated in the simulations of the FFA4 complex with TUG-891, which reduced the repulsion of the carboxyl groups of nearby E204^5.35^, D208^5.39^ and TUG-891. Whilst mutation of E204 to Ala reduced the potency of TUG-891 this alteration did not alter the potency of DHA. By contrast, whilst mutation of D208 to Ala was without effect on potency of TUG-891 this alteration significantly increased the potency of DHA (**Fig. S7a**), consistent with differences in the detailed location of the carboxyl of the synthetic and fatty acid ligands (**Fig. 5**). The receptor and agonists had small fluctuations, with DHA having a higher mobility, in all the three 1µ MD simulations (**Table S3**). From the average ligand-residue interaction energy, DHA and TUG-891 showed electrostatic attraction to nearby FFA4 residues R24^N^ and R22^N^ and repulsion to E204^5.35^ (**Fig. 5a and b**). However, mutation of either or both R24^N^ and R22^N^ residues to alanine did not alter ligand potency (**Fig. 2e, Fig. S7a**). In the simulations, in the absence of the agonists, R22^N^ and R24^N^ interact with E204^5.35^ and D208^5.39^, thus, occluding ligand access to the interhelical hydrophobic cavity. TUG-891, with its carboxyl group located deeper in the binding cavity, is further stabilized by electrostatic interactions with W198^ECL2^. The MD simulations allowed us to observe water clusters at the extracellular vestibule of the orthosteric binding site of FFA4 (**Fig. 5a and b**) which mediate polar interactions between the agonists, E204^5.35^, D208^5.39^, W198^ECL2^, R24^N^ and R22^N^. This supports the presence of water as suggested by the cryo-EM density (**Fig. 2f**).

**Figure 5.**
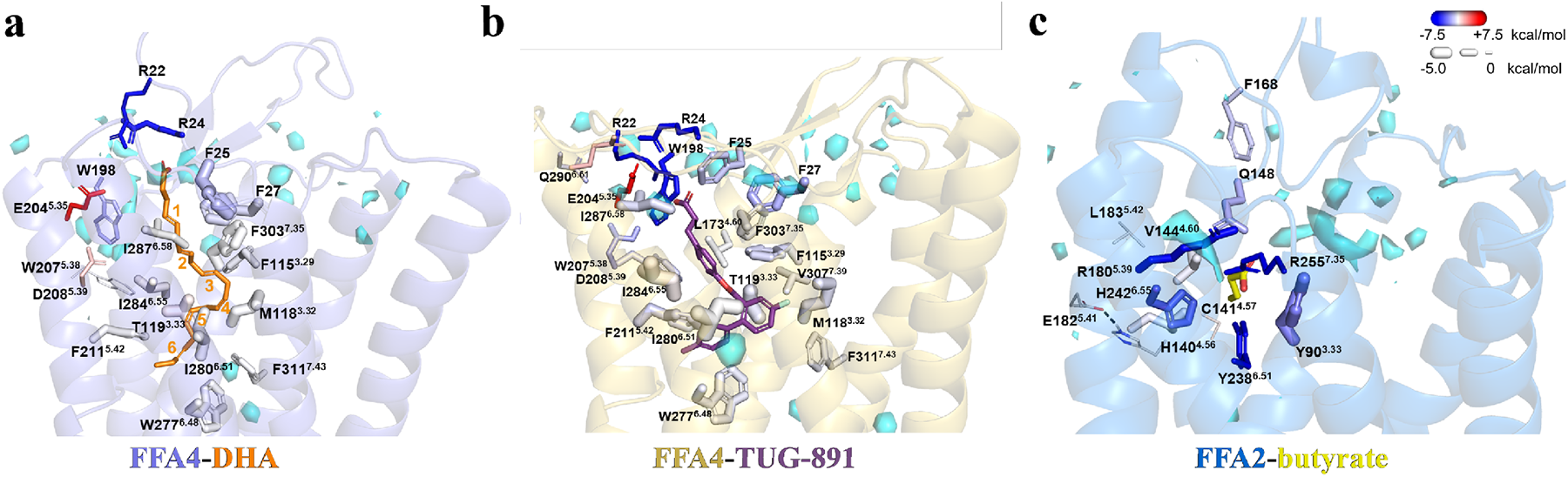
Agonist recognition of FFA4 and FFA2 probed by MD simulations. **(a)** FFA4-DHA, **(b)** FFA4-TUG-891, and **(c)** FFA2-butyrate complexes. A representative frame is shown with key residues forming contacts with DHA (orange) with DHA carbon-carbon double bonds numbered 1-6, TUG-891 (purple) and butyrate (yellow) in stick representation. The size and color of the residues correspond to the average strength of van der Waals and electrostatic interactions with the agonist, respectively. Water clusters observed in the MD simulations are shown in the cyan surface-like representation. The superscripts in the amino acid labels denote the Ballesteros–Weinstein generic GPCR residue numbering.

According to ligand fragment interaction energy calculations, the carboxyl group of the FFA4 agonists is further stabilized by electrostatic attraction to electron-deficient aromatic hydrogens of F25^N^ and F27^N^ and van der Waals interactions with their aromatic rings (**Fig. 5a, Table S4**). The first two double bonds of DHA demonstrate high mobility and do not form persistent interactions with surrounding residues. Deeper inside the pocket, double bonds 3-5 form stable van der Waals interactions with F115^3.29^, M118^3.32^, and T119^3.33^ (**Fig. 5a**). The last double bond of DHA is engaged in van der Waals interactions with F211^5.42^, I280^6.51^, and the ‘toggle switch’, W277^6.48^ (**Fig. 5a**). A similar picture is observed for TUG-891, with its last two aromatic rings, having interaction energy with these residues, along with I287^6.58^ and I284^6.55^ (**Fig. 5b**), mutation of which to Ala, as noted earlier, reduces potency of TUG-891 (**Fig. S7a**).

In FFA2, butyrate is stabilized by strong electrostatic interactions with R180^5.39^ and R255^7.35^, together with Y238^6.51^, Y90^3.33^ and Q148^ECL2^ (**Fig. 5c**), as suggested by the cryo-EM structure. In the simulations, we also observed electrostatic stabilization of the ligand by H242^6.55^ (**Fig. 5c**). The hydrogen bond between H140^4.56^ and E182^5.41^ suggested by cryo-EM was found to be persistent throughout the MD simulations.

No large movements of the helices in FFA2 and FFA4 occurred upon the removal of the agonist and miniG_q_ protein during 1µs simulations. However, we did observe the start of deactivation processes of the receptors. We saw an increase in mobility of the aromatic residues at positions 5.47, 6.44 and 6.48 (**Fig. S9**) associated with GPCR activation ^74,75^. In addition, the formation of the ‘ionic lock’ involving E^3.49^ and R^3.50^ of the ERY motif, and the conformational change of the microswitch residue at position 7.53 of the NPxxY motif, both leading to an inactive state of GPCRs, were observed in the receptors (**Fig. S10**). These changes were observed in all simulations lacking the miniG_q_ protein, usually to a greater extent in the systems without the agonist.

In summary, the MD simulations support the importance of hydrophobic and aromatic contacts deep inside the interhelical cavity in FFA4 as opposed to the polar contacts at the extracellular cavity of FFA2.

### Activation mechanisms of FFAs

As inactive structures of FFA4 have not been experimentally solved, we took an inactive structure model (FFA4-AF) obtained from the GPCRdb database ^76^, which was generated using an AlphaFold-based multi-state prediction protocol ^77^, in our structural comparison analysis. Our analysis showed significant conformational changes at the cytoplasmic region, including the outward and inward displacements of TM6 and TM7, respectively, as observed in the activation of other class A GPCRs ^75,78,79^, when comparing the active DHA-bound structure and the inactive structure of FFA4 (**Fig. 6a**). The extracellular region exhibits rather modest conformational differences except for the N-terminal region. In the FFA4-AF, the N-terminal region sticks out and does not interact with the rest of the receptor (**Fig. 6a**). This contrasts the ‘U’ shape N-terminal segment that is buried inside the 7-TM bundle in the active FFA4, which is likely stabilized by the interaction between the N-terminal residue F27 and DHA (**Fig. 1c and 6a**).

**Figure 6.**
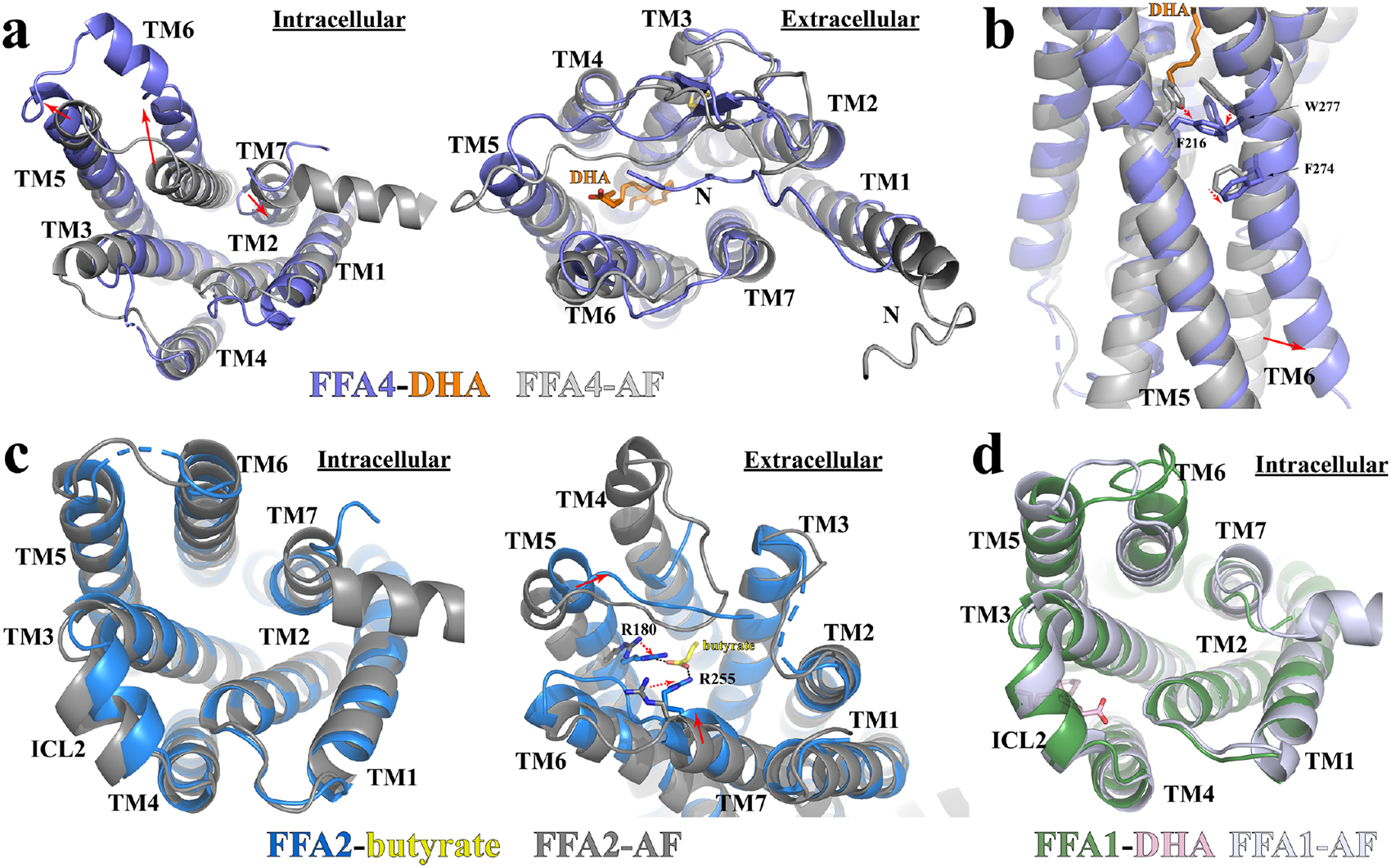
Activation of FFAs. **(a)** Superimposition of the active DHA-bound FFA4 structure (slate) to the Alphafold predicted inactive FFA4 structure FFA4-AF (light grey) viewed from the intracellular (left) and the extracellular (right) sides. **(b)** Residues involved in the receptor activation at the core region of FFA4. **(c)** Superimposition of the active butyrate-bound FFA2 structure (blue) to the Alphafold predicted FFA2 structure FFA2-AF (dark grey) viewed from the intracellular (left) and the extracellular (right) sides. **(d)** Superimposition of the active DHA-bound FFA1 structure (green) to the Alphafold predicted FFA1 structure FFA1-AF (light blue) viewed from the intracellular side. Red solid and dash arrows represent conformational changes of TMs and individual residues, respectively, from the Alphafold predicted structures to the active agonist-bound structures of FFA1, FFA2, and FFA4.

At the bottom of the ligand-binding pocket of inactive FFA4, a triad of aromatic residues F216^5.47^, F274^6.44^, and W277^6.48^ form an aromatic cluster with extensive π-π interactions (**Fig. 6b**). In the active structure of FFA4-DHA, the long chain of DHA reaches this motif and causes rearrangements of the three residues (**Fig. 6b**). In many Class A GPCRs, W^6.48^ and F^6.44^ constitute a conserved ‘activation switch’ microdomain, and conformational rearrangement of this microdomain serves as a crucial step in the activation mechanism. ^78,80–82^. Indeed, the movement of F274^6.44^ and W277^6.48^ breaks the continuous helical structure of TM6 of FFA4 (**Fig. 6b**), resulting in an outward displacement of the cytoplasmic segment of TM6, a hallmark of GPCR activation.

When we tried to compare the AlphaFold-predicted inactive structure of FFA2 obtained from the GPCRdb database (FFA2-AF) with our active structure of FFA2-butyrate, surprisingly, we found that FFA2-AF closely resembles the active conformation of FFA2 with only subtle differences at the cytoplasmic region, indicative of an active conformation of FFA2-AF (**Fig. 6c**). This finding complicates the examination of conformational changes during receptor activation. Nonetheless, we observed inward movement of TM5 and TM7 at the extracellular region in the FFA2-butyrate structure compared to the ligand-free structure of FFA2-AF, which is likely due to interactions between butyrate and the two arginine residues R180^5.39^ and R255^7.35^ (**Fig. 6c**). Similar inward movement of the extracellular segment of TM5 was also observed in the β2-adrenergic receptor (β2AR) during receptor activation due to hydrogen bonds between agonists and two serine residues of TM5, leading to the rearrangement of the P^5.50^/I^3.40^/F^6.44^ motif at the core region of the 7-TM bundle and outward movement of TM6 ^83,84^. The PIF motif functions as a molecular microswitch in the activation of some class A GPCRs ^79,85^. We hypothesize that FFA2 adopts a similar activation mechanism, where the agonist-induced inward movement of TM5 leads to the rearrangement of the core triad motif P191^5.50^/T97^3.40^/F231^6.44^ and the outward movement of TM6.

For FFA1, the AlphaFold-predicted inactive structure (FFA1-AF) also displayed subtle conformational differences compared to the active miniG_q_-coupled FFA1 in the intracellular region, suggesting that FFA1-AF may adopt an active-like conformation (**Fig. 6d**). In addition, the DHA binding site in FFA1 cannot be definitively resolved at this time, making it difficult to speculate on the mechanism of DHA-mediated activation of the receptor. If DHA binds to ‘Site 2’, it is plausible that it stabilizes the helical structure of ICL2, similar to FFA1 ago-PAMs, to position it for interaction with G protein. Furthermore, DHA interacts directly with mini-G_αq_ at ‘Site 2’ (**Fig. 3b**), indicating that it may also function to directly stabilize the interactions between FFA1 and G_q_.

### Insights into the coupling of G_q_ to FFAs

In our structural studies, we used the miniG_q_ variant of the heterotrimeric Gq due to its greater propensity to form stable complexes with FFAs in our experiments. Consistent with other GPCR-G protein complexes, the C-terminal α5 helix ^86^ of miniG_αq_ served as the primary interaction site for the receptors (**Fig. 7a**). Remarkably, the orientations of α5 of miniG_αq_ with respect to the three receptors were highly similar (**Fig. 7a**), with each receptor forming similar sets of interactions with the wavy hook region in α5 of mini-G_αq_ (**Fig. S11**). The C-terminal α5 helix of mini-G_αq_ is the main interaction site for the receptors. As mentioned previously, we did not observe strong cryo-EM density for a putative helix 8 (H8) in any of the four structures. This suggests a disordered C-terminal region following TM7 in the three active receptors. However, we did observe direct interactions between the intracellular end of TM7 of each receptor and mini-G_αq_ (**Fig. S11**), underscoring an important role of TM7 in G_q_-coupling to each of the three receptors.

**Figure 7.**
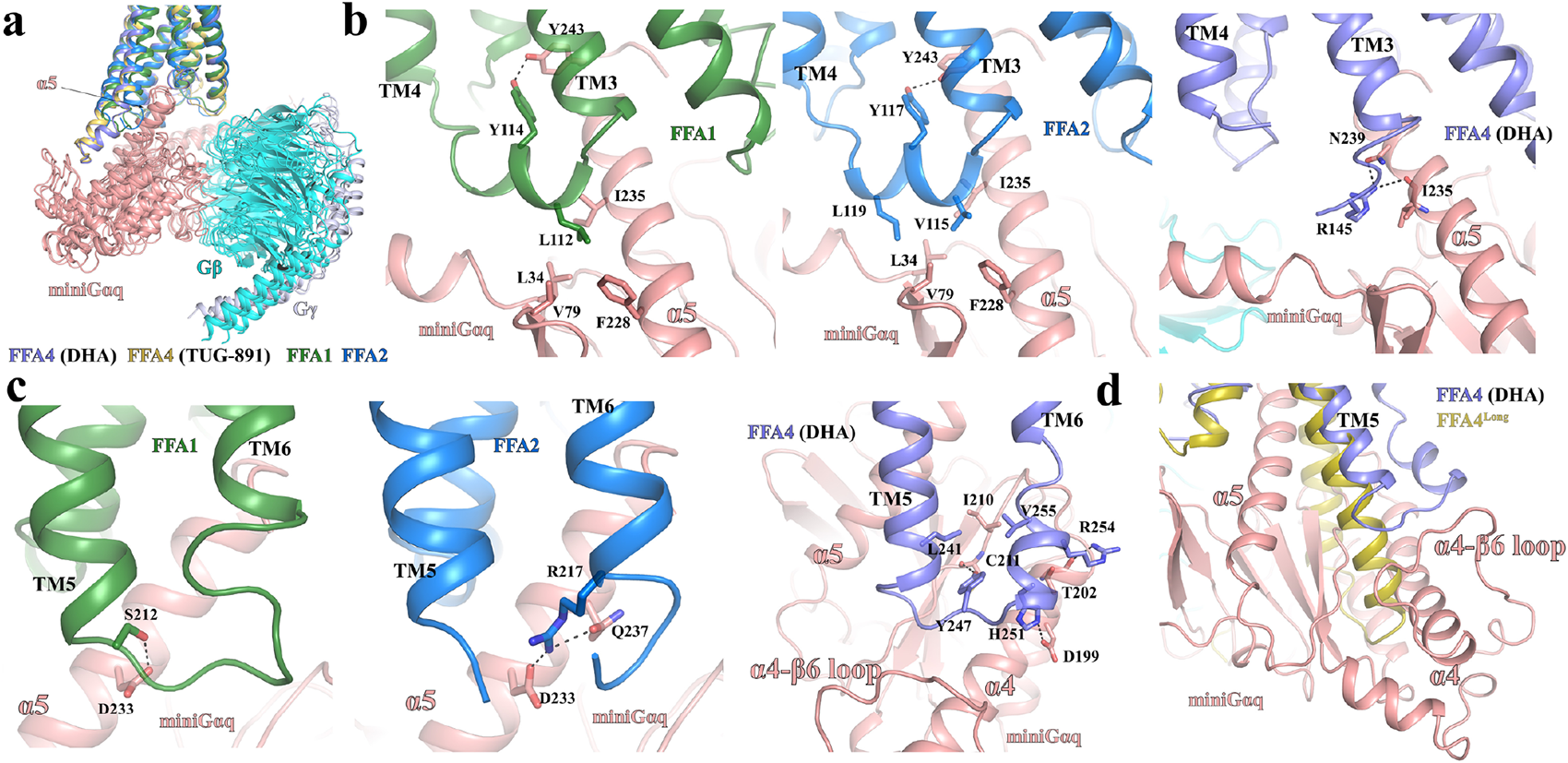
Differences in the coupling of miniG_q_ to FFAs. **(a)** Alignment of the structures of FFA1, FFA2, and FFA4 coupled with miniG_q_ based on the receptors. **(b)** Differences in the interactions between miniG_αq_ and ICL2 of FFA1, FFA2, and FFA4. **(c)** Differences in the interactions between miniG_αq_ and ICL3 of FFA1, FFA2, and FFA4. **(d)** Superimposition of the AlphaFold predicted structure of FFA4^Long^ to the structure of DHA-bound FFA4 coupled with miniG_q_. MiniG_αq_, G_β_ and G_γ_ subunits are colored in salmon, cyan and light blue, respectively. The colors of receptors and ligands are indicated in each panel.

Although the interactions at the α5 helix of miniG_αq_ are similar across all three receptors, they engage in distinct interactions with other regions of miniG_q_. Specifically, for FFA1 and FFA2, the intracellular loop 2 (ICL2) forms a helical structure to directly interact with miniG_q._ In both receptors, a conserved tyrosine residue, Y114^ICL2^ in FFA1 and Y117^ICL2^ in FFA2, sticks toward the 7-TM core and forms a hydrogen bond with Y243 of mini-G_αq_ (**Fig. 7b**). Additionally, on the opposite side of ICL2, V115 ^ICL2^ and L119 ^ICL2^ in FFA2, or L112 ^ICL2^ in FFA1, form hydrophobic interactions with mini-G_αq_ residues L34, V79, F228, and I235 (**Fig. 7b**). A similar set of hydrophobic interactions are also observed in the structures of the muscarinic acetylcholine receptor 1 (M1R)-G_11_ ^87^ and the serotonin receptor 5-HT_2A_-miniG_q_ complexes ^47^. However, in FFA4, a large part of ICL2 is disordered, and no similar hydrophobic interactions are observed (**Fig. 1b**). Nevertheless, R145^ICL2^ of FFA4 forms hydrogen bonds with the side chain of N239 and the main chain carbonyl of I235 in the α5 helix of mini-G_αq._

ICL3 is another region in the three receptors that interacts differently with miniG_q_. In FFA1 and FFA2, ICL3 is positioned close to the wavy hook region of mini-G_αq_ (**Fig. 7c**). S212^ICL3^ in FFA1 forms a hydrogen bond with D233 of miniG_αq_, while R217^ICL3^ in FFA2 forms hydrogen bonds with D233 and Q237 of miniG_αq_ (**Fig. 7c**). However, in FFA4, TM5 extends by two helical turns compared to that of FFA1 and FFA2, resulting in ICL3 of FFA4 being positioned near the α4 helix and the α4-β6 loop of mini-G_αq_ (**Fig. 7c**). In this position, a segment of ICL3 in FFA4 adopts a helical structure. In the structure of FFA4-miniG_q_ with DHA, residues Y247^ICL3^ and R254^ICL3^ of FFA4 form hydrogen bonds with the carbonyl groups of the mini-Gαq residues C211 and T202 backbone, respectively, while H251^ICL3^ of FFA4 forms a hydrogen bond with D199 of mini-G_αq_ (**Fig. 7c**). Additionally, V255^ICL3^ and L241^5.72^ of FFA4 pack against I210 of mini-G_αq_ to form hydrophobic interactions (**Fig. 7c**).

The conformation of TM5 and ICL3 may account for the inability of FFA4^Long^, the long form of FFA4, to induce G_q/11_ signaling ^43,61^. FFA4^Long^ contains an insertion of 16 additional amino acids after Q232^5.63^, which would further extend TM5 and potentially cause a severe steric clash with G_αq_. This clash is clearly visible if we superimpose the AlphaFold-predicted structure of FFA4^Long^ onto FFA4 in our structure (**Fig. 7d**).

## Discussion

### Lipid recognition by FFAs

The different binding pockets in FFA2 and FFA4 for their endogenous SCFA and LCFA ligands clearly explain their preferences of free fatty acids ^39,40^. For FFA2, the small size of the binding pocket can only accommodate LCFAs with very short carbon chains. For FFA4, the binding pocket is much larger to accommodate LCFAs. It seems that the binding of LCFAs to FFA4 is largely driven by hydrophobic and π-π interactions since the mutations of polar residues near the carboxylate group of DHA did not significantly reduce the potency. SCFAs and medium-chain fatty acids (MCFAs) with shorter carbon chains would result in less contacts with FFA4 and thus weaker potency for this receptor. On the other hand, despite the small observed binding pocket of FFA2, a number of larger synthetic FFA2 selective and orthosteric agonists have been identified and studied ^70,88^. It will be interesting in time to explore how FFA2 recognize those large ligands.

The molecular basis for the lipid recognition of FFA1 is not readily clear based on our structural data. ‘Site 1’ and ‘Site 2’ are both potential binding sites for DHA. ‘Site 1’ but not ‘Site 2’ is rich in aromatic residues, which would favor the binding of PUFAs to FFA1. In addition, the two arginine residues R158^5.39^ and R258^7.35^ that are highly conserved in FFA1-3 form salt bridges with the carboxylate group of DHA (**Fig. 3a**). Mutations of these two residues significantly reduced the potency of DHA and another LCFA, γ-linoleic acid (γ-LA) ^65^, further suggesting that ‘Site 1’ is the primary binding site for DHA. However, the strong cryo-EM density in ‘Site 2’ in our structure implies potential binding of DHA at this site as well, which may serve as the secondary binding site for DHA. Nevertheless, neither ‘Site 1’ nor ‘Site 2’ is large enough to accommodate the entire LCFAs with 20 carbons or more. The mechanism for the selectivity of FFA1 for LCFAs over SCFAs still needs further investigation.

### Insight into drug development on FFAs

Over the past decade, a growing body of research has established the critical roles of FFAs in regulating metabolism and immunity. Studies have also provided evidence suggesting that agonists of FFA1, FFA2, and FFA4 have the potential to treat metabolic and inflammatory diseases. However, only the FFA1 agonist TAK-875 has completed all three phases of clinical trials. While TAK-875 showed promising results in improving glycaemic control with a low risk of hypoglycemia in a phase III trial for diabetics, its further development was halted due to liver toxicity concerns ^89^. The underlying mechanism behind TAK-875’s liver toxicity is still not entirely clear, with studies suggesting both direct hepatoxicity in an FFA1-dependent manner and metabolite-induced inhibition of hepatic transporters and mitochondrial respiration ^90,91^. To advance future drug development, it is crucial to determine whether TAK-875’s liver toxicity is related to the activation of FFA1. The two well-defined ligand binding sites in FFA1 offer opportunities for designing chemically diverse FFA1 ligands. If TAK-875’s liver toxicity is caused by its metabolites, developing new FFA1 agonists with novel chemical scaffolds may provide a solution to this issue.

Compared to FFA1, the development of synthetic agonists for FFA2 and FFA4 toward the clinic has been limited, with only a small number of FFA2 ligands having been reported ^39,70,88,92^. By contrast, although the chemical diversity of synthetic FFA4 agonists has been rather limited ^22^, many analogs of the TUG-891 phenyl-propionic scaffold have been generated to improve the drug-like characteristics of this ligand for further assessment of their effects on the regulation of glucose homeostasis and other disease indications. A surprising feature of the observed binding of TUG-891 and DHA within FFA4 was the absence of interaction between the carboxylate of agonists and R99^2.64^. This had been widely anticipated based on earlier modelling and mutagenesis studies. Initial modelling studies linked to the development of TUG-891 predicted an ionic interaction with this residue ^48,93^ and subsequent mutagenesis to R99Q confirmed the importance of this residue as the agonists were unable to activate this mutant. However, R99^2.64^ does not interact directly with agonists but rather forms polar interactions with D30^N^ and E43^1.35^ and a cation-π interaction with F304^7.36^ (**Fig. S12**). By doing so, R99^2.64^ likely stabilizes the aromatic network of F25^N^, F27^N^, F28^N^, F115^3.29^ and F303^7.35^ that forms a lid to the hydrophobic pocket. Mutation of R99^2.64^, therefore, could break this aromatic network and lead to the exposure of the hydrophobic pocket to water destabilizing agonist binding.

Our work and another recent structural study ^61^ show that the carboxyl group of ligands does not form specific interactions with FFA4; rather binding and ligand location is driven through hydrophobic interactions. The observed orthosteric binding pocket of FFA4, as revealed by our structures, offers valuable templates for designing new agonists for this receptor using computer-aided and AI-driven drug design (CADD and AIDD) approaches. Another group of FFA4 agonists indeed don’t contain the carboxyl group but have a sulphonamide or amide moiety as the central core linking the aromatic rings to form a ‘L’ shape ^22,39^. Interestingly, when docking one such ligand, TUG-1197, into our FFA4 structure, the sulphonamide formed an H-bond with T119^3.33^ and its position overlapped with the O-linker of TUG-891 (**Fig. 2d**). It is hence noteworthy that TUG-1197 displayed markedly reduced potency at a T119A mutant of FFA4, and although more modest in extent the potency of TUG-891 was also reduced at T119A.

In contrast to FFA4, key residues of the orthosteric binding pocket of FFA2 were highly aligned with those predicted by previous mutagenesis studies ^68^. Given the challenges of developing potent and selective orthosteric FFA2 activators, there has been interest in the availability and design of selective allosteric agonists of FFA2 ^22,39^. Although nothing is currently known about their mode of binding, structural studies akin to those reported herein, and the ability to use computational tools to predict allosteric binding sites ^94^, it is likely that rapid progress could be made. This will also allow the development of more ‘drug-like’ allosteric regulators of FFA2 and potentially also of the other SCFA receptor FFA3 where no high potency synthetic ligands are currently available ^39^, even as tool compounds, to better explore the biology and potential patho-physiological functions of this receptor. In both FFA2 and FFA4, we observed strong cryo-EM density in sites similar to the ‘Site 2’ observed in FFA1, where we modeled PI4P (phosphatidylinositol-4-phosphate) and palmitic acid to fit the density (**Fig. S6**). These observations suggest the possibility of developing allosteric modulators for FFA2 and FFA4 targeting this site. In the case of FFA2, we also observed a large cavity at the extracellular region right above the butyrate binding pocket. This cavity represents another potential allosteric site, which we refer to as ‘Site 3’. Interestingly, if the structure of FFA2 is aligned with the structure of the muscarinic receptor M2R bound to a positive allosteric modulator (PAM) named LY2119620, the allosteric site of LY2119620 highly overlaps with the putative ‘Site 3’ in FFA2. As noted earlier there are a number of FFA2 PAMs reported ^40^. They may target the two potential allosteric sites, ‘Site 2’ and ‘Site 3’, revealed by our structures. Further structural studies are needed to fully understand the molecular mechanisms of allosteric modulation in FFAs.

## Acknowledgement

We thank the cryo-EM facility at the University of Pittsburgh partly supported by the grants S10 OD025009 (Krios) and S10 OD019995 (Falcon 2/3 camera) from the National Institutes of Health (NIH) in the USA. We thank Dr. Sudha Chakrapani for oversight of the cryo-EM core facility at Case Western Reserve University. This work was supported by the NIH grant R35GM128641 to C.Z., the Medical Research Council (U.K.) grant MR/X010198/1 to G.M., the Biotechnology and Biological Sciences Research Council (U.K.) grants BB/R001480/1 and BB/S000453/1 to G.M. and BB/R007101/1 to I.G.T..

This project made use of computational time on Kelvin-2 supported by Engineering and Physical Sciences Research Council (EPSRC) (grant no. EP/T022175/1 and EP/W03204X/1) and ARCHER2 granted via the UK High-End Computing Consortium for Biomolecular Simulation, HECBioSim (https://www.hecbiosim.ac.uk), supported by EPSRC (grant no. EP/R029407/1 and EP/W03204X/1).

## Contributions

C.Z., G.M., and I.G.T. conceived the project and designed the research with X.Z. X.Z. performed protein expression and purification studies, screened cryo-EM grids, collected cryo-EM data, and processed the data under the supervision of C.Z. L.J. performed pharmacological and mutagenesis studies, A-A.G. performed computational analysis. G.M. and I.G.T. analyzed data. K.L. assisted in the cryo-EM data collection. C.Z. wrote the manuscript together with G.M. and I.G.T. with the help from X.Z.

## Data availability

The 3D cryo-EM density maps of FFA signaling complexes have been deposited in the Electron Microscopy Data Bank under the accession numbers EMD-41013 and EMD-41014 for the DHA-FFA1-miniGq complex, EMD-41010 for the butyrate-FFA2-miniGq complex, and EMD-41007 and EMD-41008 for the DHA- and TUG-891-FFA4-miniGq complexes, respectively. Atomic coordinates for the atomic models have been deposited in the Protein Data Bank (PDB) under the accession numbers 8T3V, 8T3S, 8T3Q, and 8T3Q for the DHA-FFA1-miniGq complex, the butyrate-FFA2-miniGq complex, and the DHA- and TUG-891-FFA4-miniGq complexes, respectively.

## Methods

### Protein expression and purification

Human FFA1, FFA2, and FFA4 were cloned into the pFastBac vector (Thermo Fisher) with the LargeBit protein ^54^ fused to the C-terminus of each receptor. The miniGαq subunit ^49^ was cloned into the pFastBac vector. Human Gβ_1_ was fused with an N-terminal His_6_-tag and a C-terminal HiBiT subunit connected with a 15-amino acid linker, which was cloned into pFastBac dual vector (Thermo Fisher) together with human Gγ_2_.

The scFv16 was expressed in High Five cells using Bac-to-Bac expression system. To purify the protein, the cell supernatant was collected and loaded onto Ni-NTA resins. Following nickel affinity chromatography, the protein was further purified by size exclusion chromatography using a Superdex 200 Increase 100/300 GL column (GE Healthcare). The purified scFv16 were pooled, concentrated and stored at -80°C until use.

Free fatty acid receptors, miniGαq and Gβ_1_γ_2_ were co-expressed in Sf9 insect cells using Bac-to-Bac method. Sf9 cells were infected with three types of viruses at the ratio of 1:1:1 for 48 h at 27 °C. After being cultured for 48 hours, the cells were harvested and frozen at -80 °C for further protein purification. Cell pellets were thawed in lysis buffer containing 20 mM HEPES, pH7.5, 50 mM NaCl, 10 mM MgCl_2_, 5 mM CaCl_2,_ 2.5 μg/ml leupeptin, 300 μg/ml benzamidine. The complexes of DHA-bound or TUG-891-bound FFA4 with miniG_q_ were assembled on cell membranes by the addition of 10 μM DHA or 10 μM TUG-891. For the FFA1-miniG_q_ and FFA2-miniG_q_ complexes, 10 μM DHA and 1 mM butyrate were added to stimulate the formation of signaling complexes. To facilitate complex formation, 25 mU/ml Apyrase (NEB), and 100 μM TCEP was added and incubated at room temperature for 2 h. The cell membranes were isolated by centrifugation at 25,000 g for 40 min and then resuspended in solubilization buffer containing 20 mM HEPES, pH7.5, 100 mM NaCl, 0.5% (w/v) lauryl maltose neopentylglycol (LMNG, Anatrace), 0.1% (w/v) cholesteryl hemisuccinate (CHS, Anatrace), 10% (v/v) glycerol, 10 mM MgCl_2_, 5 mM CaCl_2_, 12.5 mU/ml Apyrase, 10 μM or 1 mM ligands, 2.5 μg/ml leupeptin, 300 μg/ml benzamidine, 100 µM TECP at 4 °C for 2 h. The supernatant was collected by centrifugation at 25,000 g for 1 h and incubated with nickel Sepharose resin (GE Healthcare) at 4 °C overnight. The resin was washed with a buffer A containing 20 mM HEPES, pH 7.5, 100 mM NaCl, 0.05% (w/v) LMNG, 0.01% (w/v) CHS, 20 mM imidazole, and 10 μM or 1 mM ligands, 2.5 μg/ml leupeptin, 300 μg/ml benzamidine, 100 µM TECP. The complex was eluted with buffer A containing 400 mM imidazole. The eluate was supplemented with 2mM CaCl_2_ and loaded onto anti-Flag M1 antibody resin. After wash, the complex was eluted in buffer A containing 5 mM EDTA and 200 μg/ml FLAG peptide and concentrated using an Amicon Ultra Centrifugal Filter (MWCO, 100 kDa). Finally, a 1.3 molar excess of scFv16 was added to the elution. The sample was then loaded onto a Superdex 200 Increase 10/300 column (GE Healthcare) pre-equilibrated with buffer containing 20 mM HEPES pH 7.5, 100 mM NaCl, 0.00075% (w/v) LMNG, 0.00025% (w/v) GDN, 0.00015% (w/v) CHS, 10 μM or 1 mM ligands and 100 µM TECP. Peak fractions of the complex were collected and concentrated to 5-10 mg/ml for cryo-EM experiments.

### Cryo-EM sample preparation and data acquisition

For cryo-EM grids preparation of DHA-FFA1-miniGq complex, butyrate-FFA2-miniGq complex and TUG-891-FFA4-miniGq complex, 3 μl of the protein complex was applied onto 300 mesh R1.2/1.3 UltrAuFoil Holey gold support films (Quantifoil). For cryo-EM grids preparation of DHA-FFA4-miniGq complex, 3 μl of the purified complex was applied to glow-discharged holey carbon grids (Quantifoil, Au300 R1.2/1.3). Grids were plunge-frozen in liquid ethane using Vitrobot Mark IV (Thermo Fischer Scientific).

All cryo-EM data were collected using Titan Krios transmission electron microscope, equipped with a Gatan K3 Summit direct electron detector and an energy filter. For TUG-891-FFA4-miniGq complex and DHA-FFA4-miniGq complex, cryo-EM movie stacks were recorded with a nominal defocus setting in the range of -1.0 to -1.8 μm using SerialEM ^95^ with beam-tilt image-shift data collection strategy with a 3 × 3 pattern and 3 shot per hole. A total of 4,968 movies for the dataset of TUG-891-FFA4-miniGq complex and 10,040 movies for three datasets of DHA-FFA4-miniGq complex were collected in the correlated double sampling (CDS) super-resolution mode of the K3 camera at a nominal magnification of 105,000× yielding a physical pixel size of 0.828 Å. Each stack was dose-fractionated to 52 frames with a total dose of 55 e^-^/Å^2^. For DHA-FFA1-miniGq complex and butyrate-FFA2-miniGq complex, 12,349 movies and 15,371 movies were collected with a nominal magnification of 105,000× using the SerialEM software running a 3 × 3 image shift pattern and 3 shot per hole. Micrographs were recorded using a super-resolution mode at a calibrated pixel size of 0.826 Å and a defocus range of -0.8 to -2.5 μm. Each stack was dose-fractionated to 50 frames with a total dose of 61.6 e^-^/Å^2^.

### Data processing, 3D reconstruction and modeling building

Image stacks were subjected to patch motion correction using cryoSPARC ^96^. The contrast transfer function (CTF) parameters were calculated using the patch CTF estimation tool in cryoSPARC.

For the TUG-891-FFA4-miniGq complex, a total of 4,928,436 particles were auto-picked and then subjected to 2D classification to discard poorly defined particles. After ab initio reconstruction and heterogeneous refinement, 391,203 particles were subjected to non-uniform refinement and local refinement, which generated a map with an indicated global resolution of 3.06 Å at a Fourier shell correlation (FSC) of 0.143. For the DHA-FFA4-miniGq complex, a threshold of CTF fit resolution of more than 4 Å was used to exclude low-quality micrographs. Each dataset was processed separately with autopicking and 2D classification. After ab initio reconstruction and heterogeneous refinement, 380,284 particles were subjected to non-uniform refinement and local refinement, which generated a map with an indicated global resolution of 3.14 Å at a Fourier shell correlation (FSC) of 0.143. For the DHA-FFA1-miniGq complex and the butyrate-FFA2-miniGq complex, a threshold of CTF fit resolution of more than 4 Å was used to exclude low-quality micrographs, respectively. A total of 7,942,319 particles and 12,559,721 particles were auto-picked and then subjected to 2D classification to discard bad particles. After ab initio reconstruction and heterogeneous refinement, 305,318 particles and 393,952 particles were subjected to non-uniform refinement and local refinement, which generated a map with an indicated global resolution of 3.39 Å and 3.07 Å at a Fourier shell correlation (FSC) of 0.143. Local resolutions of density maps were estimated in cryoSPARC.

The Alphafold-predicted structures of FFA1, FFA2, and FFA4 served as initial models for model rebuilding and refinement against the electron microscopy map. The model was initially docked into the electron microscopy density map using Chimera ^97^. This step was followed by iterative manual adjustment and rebuilding in COOT ^98^. Real space refinement and Rosetta refinement were then carried out using Phenix programs ^99^. To validate the model, MolProbity was employed for assessing its structural statistics ^100^.

For the preparation of structural figures, Chimera and PyMOL (https://pymol.org/2/) were utilized. The final refinement statistics can be found in Supplementary Table 1. To evaluate the degree of overfitting during the refinement process, the final model was refined against one of the half-maps. The resulting map versus model FSC curves were compared with both half-maps and the full model. Surface coloring of the density map was achieved using UCSF Chimera ^97^.

### Mutagenesis and bioluminescence resonance energy transfer (BRET)-based arrestin-3 recruitment assays

All cell culture reagents and TMB substrate solution were from Thermo Fisher Scientific (Loughborough, UK). Polyethylenimine (PEI) [linearpoly(vinyl alcohol) (MW-25000)] was from Polysciences (Warrington, PA). Molecular biology enzymes and reagents were from Promega (Southampton, UK). TUG-891 and docosahexaenoic acid (DHA) were from Tocris Biosciences (Bristol, UK). TUG-1197 was a gift from Trond Ulven (University of Copenhagen).

A plasmid encoding the short isoform of human FFA4 fused at its C terminus to enhanced yellow fluorescent protein (eYFP) and containing an N-terminal FLAG epitope tag was generated as described previously ^101^. Mutations were introduced into the FFA4 sequence using the QuikChange method (Stratagene), and in all cases the presence of the mutation was verified through sequencing. The NanoLuc luciferase coding sequence was subcloned after PCR amplification (using primers designed to add Xba1 and NotI sites) into an arrestin-3-pcDNA3 plasmid (arrestin-3-NLuc).

All FFA4 constructs were expressed in HEK293T cells, which were maintained in Dulbecco’s modified Eagle’s medium supplemented with 0.292 g/l L-glutamine, 1% penicillin/streptomycin mixture, and 10% heat-inactivated fetal bovine serum at 37 °C in a 5% CO_2_ humidified atmosphere. To express receptors, cells were transfected using PEI. The day before transfection 2 × 10^6^ cells were plated into 10 cm dishes. Plasmid DNA was then combined with PEI (in 1:6 ratio) in 500 μl of 150 mM NaCl, thoroughly mixed then incubated for 10 min at room temperature. Cell medium was changed and the DNA–PEI mixture was added to the medium in a dropwise manner.

For BRET assays, HEK293T cells were seeded in 10 cm^2^ dishes and transiently co-transfected with wild type or each of the indicated FFA4 mutants, each with a FLAG epitope tag engineered into the N-terminal domain and eYFP fused at its C terminus, and arrestin-3-NLuc at a 50:1 ratio respectively using PEI. Control cells were transfected with arrestin-3-NLuc only. After 24 h cells were detached by incubating with trypsin-EDTA and seeded at 5 x 10^4^ cells/well in poly-D-lysine coated white 96-well plates, then incubated overnight at 37°C. Cells were washed once with pre-warmed (37°C) HBSS and incubated in HBSS for 30–60 min at 37°C. The luciferase substrate coelenterazine-h (Prolume) was added to a final concentration of 5 μM and the plate incubated for 10 min at 37°C protected from light.

Agonists were added at the relevant concentrations in triplicate and the plate incubated for a further 5 min at 37°C, then the emissions at 475 nm and 535 nm were read on a PHERAstar FS. The net BRET ratio (mBRET) was calculated as follows: [(signal 535 nm/signal 475 nm) – (signal nanoluc luciferase only 535 nm/signal nanoluc luciferase only 475 nm)] *1000. Direct measures of e-YFP fluorescence determined total expression levels of FFA4 receptor variants.

### Cell surface enzyme-linked immunosorbent assays

Cell surface expression of receptors was quantified by live-cell ELISA. The same co-transfected cells used for BRET studies were seeded at 5 x 10^4^ cells/well in poly-D-lysine-coated clear 96-well plates and incubated overnight at 37°C. Cells were incubated with primary antibody (rabbit polyclonal anti-FLAG 1:1000) in culture medium for 1 h at 37°C, then washed three times with DMEM-HEPES and incubated with secondary antibody (Horseradish peroxidase (HRP)-sheep anti-rabbit IgG 1:10,000) for 1 h at 37°C protected from light. Cells were then washed three times with warmed (37°C) PBS. Finally, PBS was removed and 100 μL/well room temperature TMB substrate was added. The plate was incubated for 15 min at room temperature protected from light, then the absorbance at 620 nm was read on a POLARStar® Omega.

### Phylogenetic analysis

Phylogenetic analysis of full-length sequences of human class A GPCRs (312 in total) was done using the sequences and tools provided by GPCRdb web server ^102^. Unrooted phylogenetic tree with increasing node order was built using FigTree v1.4.4 ^103^. All GPCRs were colored according to their GPCRdb ligand type. Nodes with descendants sharing the same GPCRdb ligand type were colored accordingly, with the remaining nodes colored grey.

### Molecular Docking

The protein structures were prepared with the protein preparation module, and the structure of TUG-1197 was assessed with the ligand preparation module of Schrodinger software. TUG-891 from the FFA4-TUG-891 complex were selected to center the docking box. Receptor docking grids with the receptor van der Waals radius scaling of 1.0 were generated. Docking poses were obtained and evaluated with the Glide program ^104–107^. The OPLS_2005 force field was used in all calculations.

### MD simulations

The cryo-EM structures of FFA4 bound to TUG-891 or DHA, and FFA2 bound to butyrate were prepared using Schrodinger Maestro 2021-3 ^108^. All molecules except the GPCR and, if necessary, the ligand and G-protein, were removed. Missing loops and sidechain atoms of the proteins were filled using knowledge-based homology modelling of the Prime module ^109–111^ with amino acid sequences taken from the UniProt database ^112^. The N- and C-termini of the receptors were extended by up to 3 residues as per the UniProt sequence, and the added residues were minimized using the 3D builder of Maestro. The N- and C-termini of the receptors and the G-protein were capped with acetyl and N-methyl groups, respectively. The obtained structures were analyzed using the Maestro Protein Reports tool and strong steric clashes were removed by local geometry minimization.

The protonation states of amino acids at pH 7.4 were predicted by PROPKA 3 as a separate application ^113,114^ and as implemented in the Maestro Protein Preparation workflow. Thus, D208^5.39^ in the FFA4-TUG-891 complex and H140^4.56^ in the FFA2-butyrate complex were kept protonated in the simulations. All non-protonated histidine residues were taken as a δ-tautomer except H242^6.55^ in the FFA2-butyrate complex, which was taken as an ε-tautomer as it reduced the root mean square deviation of the ligand atoms.

The input for membrane simulations was prepared using the CHARMM-GUI server ^115–123^. The receptor was oriented in membrane using the PPM server within CHARMM-GUI ^124^. The ligand parameter files were created by the Antechamber utility of the CHARMM-GUI server with the AM1BCC charge scheme and GAFF2 atom types. The receptor was placed in the 1-palmitoyl-2-oleoyl-sn-glycero-3-phosphocholine (POPC) bilayer membrane sized 100 Å x 100 Å and 22.5 Å solvent layer on each side of the membrane. For receptor-miniGq systems, 120 Å x 120 Å membrane bilayer was used. The solvent contained 150 mM NaCl, and the total number of atoms was between 100,000 and 130,000 atoms for receptor-only systems and between 240,000 and 260,000 atoms for the receptor-miniGq systems.

The minimization, equilibration and production were done using the PMEMD program from the Amber20 package [AMBER 20] using the ff19SB ^125^, LIPID21 ^126^, GAFF2 ^127^ force fields and the TIP3P model for the protein, membrane lipids, ligands and water, respectively. The nonbonded interaction cut-off was set as 9 Å.

As per the recommended CHARMM-GUI protocol, the initial energy minimization included 5000 steepest descent steps followed by 5000 more steps using the conjugated gradient method. The equilibration steps followed the same pattern of gradually decreasing the force constants of positional restraints but were conducted with longer simulation length than recommended by the CHARMM-GUI protocol. Heating to 310 K was done in the NVT ensemble with 1 fs timestep in two consecutive equilibration steps, each 2.5 ns long. This was followed by a 2.5 ns NPT equilibration with Berendsen barostat and a 1 fs timestep ^128^. The next two steps of NPT equilibration used 2 fs timestep and lasted for 10 ns each, and the final step in which only the protein backbone was restrained lasted 20 ns. The production was done with Langevin thermostat ^129^ with a friction coefficient of 1.0 ps^−1^ and Berendsen barostat with timestep of 2 fs. The production was done in 3 replicas of 1 µs for the empty and ligand-bound receptor systems and 1 replica of 1 µs for the ligand bound receptor-miniGq systems. The snapshots of the simulations were saved every 50’000 steps, or 0.1 ns, in simulations.

The results of simulations were analysed using CPPTRAJ from the Amber20 package and MDAnalysis ^130,131^. The residue–ligand interaction energy was calculated using the “namdenergy.tcl” script v1.6 for VMD ^132^ and the NAMD2 program with cut-off and switch parameters of 9 Å and 7.5 Å, respectively ^133,134^. Forcefield parameters were taken from the AMBER parameter file used for simulations. Energy calculation was done for every 10^th^ snapshot, or every 1.0 ns of simulation.

## Supplemental figures

**Figure S1.**
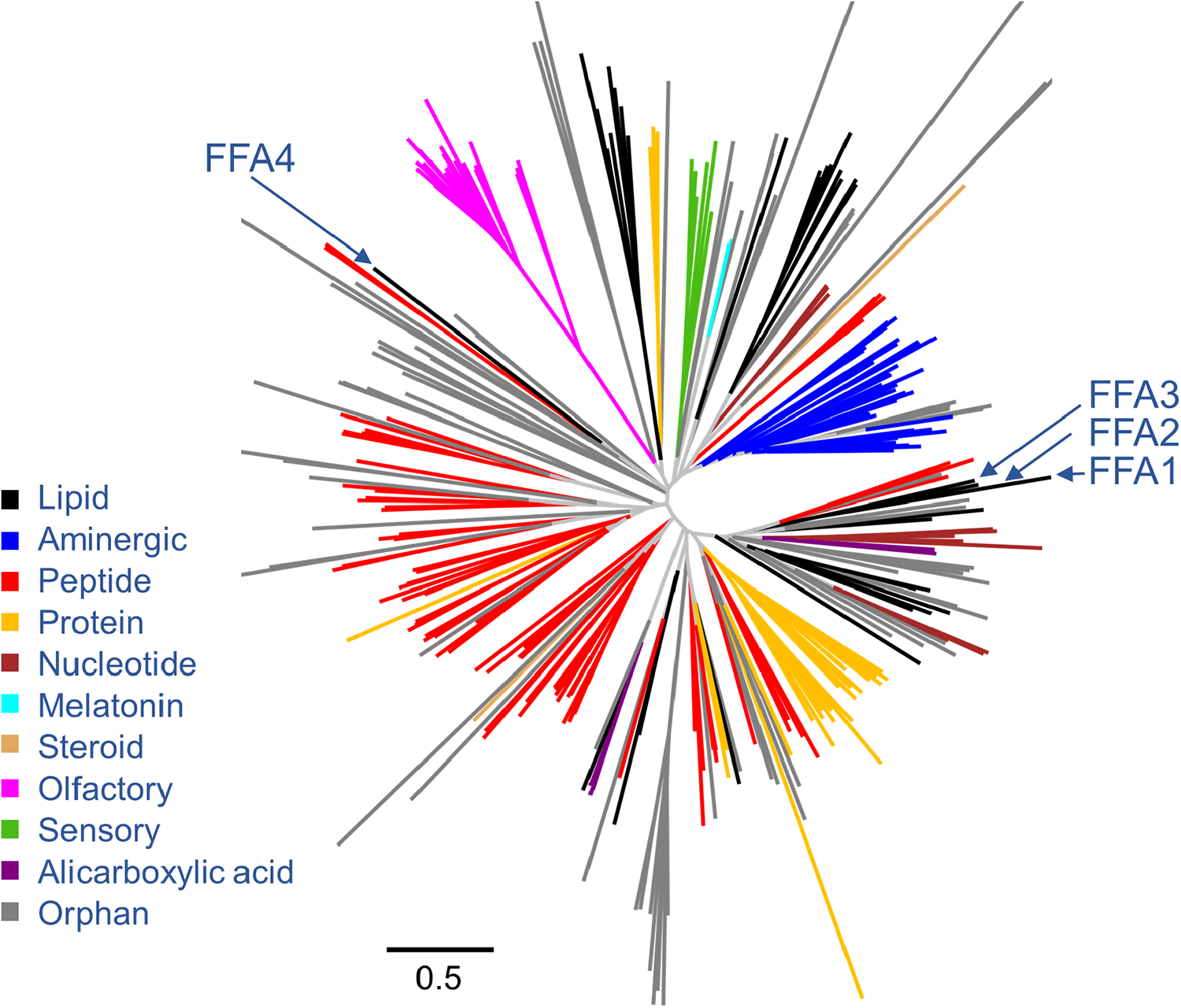
Phylogenetic analysis of the FFA receptor family. Clustering of full-length sequences of human class A GPCRs (312 in total) was done using the sequences and tools provided by the GPCRdb web server (https://gpcrdb.org/). An unrooted phylogenetic tree with an increasing node order was built using FigTree v1.4.4(http://tree.bio.ed.ac.uk/). All GPCRs were colored according to their GPCRdb ligand type. Nodes with descendants sharing the same GPCRdb ligand type were colored accordingly, with the remaining nodes colored grey.

**Figure S2.**
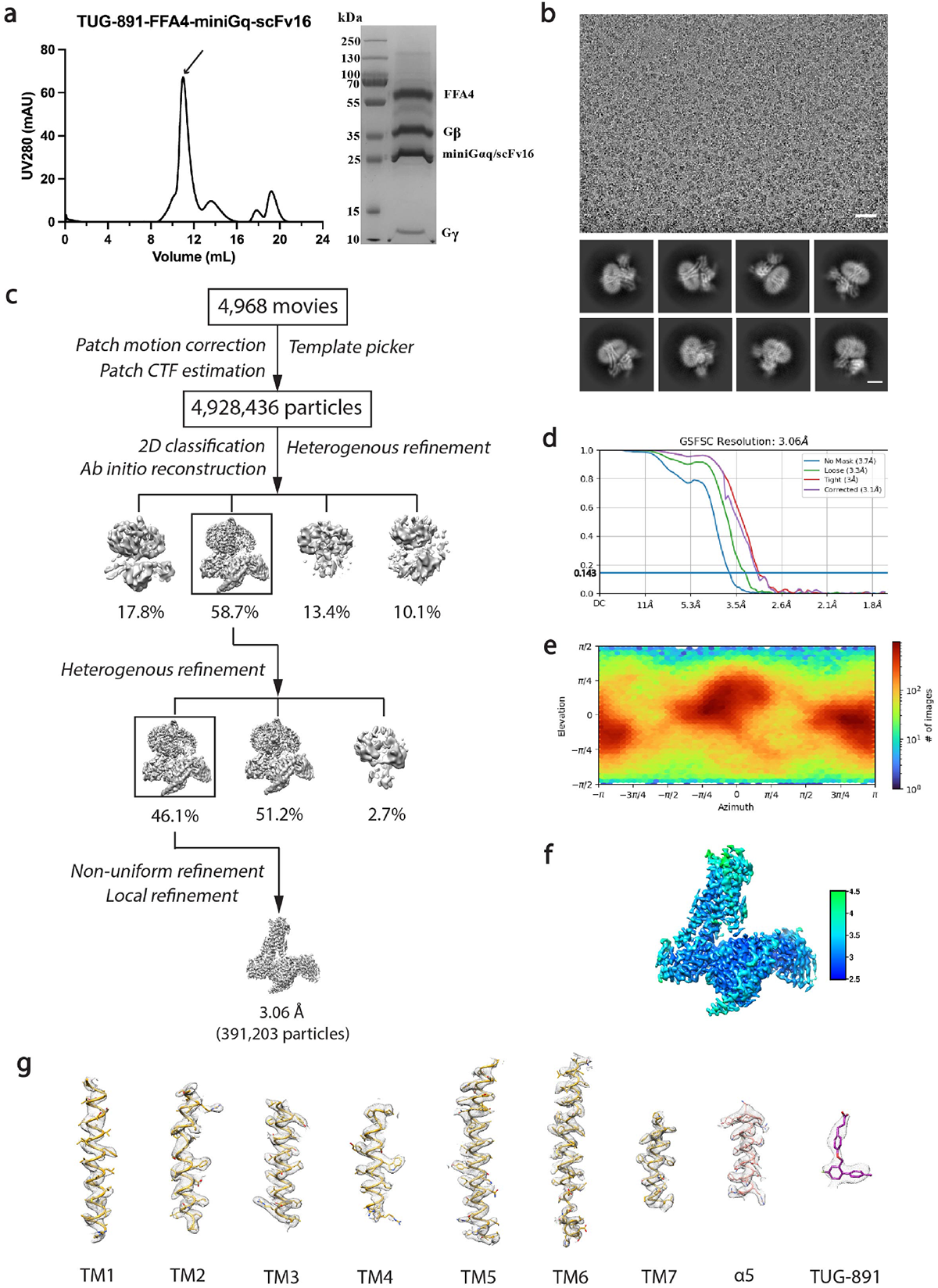
Purification of the FFA4-miniGq complex with TUG-891 and cryo-EM data processing. **(a)** Size-exclusion chromatography profile and SDS-PAGE analysis of the purified TUG-891-FFA4-miniGq complex. **(b)** Representative cryo-EM micrograph (scale bar: 50 nm) and 2D class averages (scale bar: 5 nm). **(c)** Cryo-EM image processing workflow for the TUG-891-FFA4-miniGq complex. **(d)** Gold-standard Fourier shell correlation (FSC) curve showing an overall resolution is 3.06 Å at FSC=0.143. **(e)** Angular distribution of the particles used in the final reconstruction. **(f)** Density map according to local resolution estimation. **(g)** Cryo-EM density maps and models of the seven transmembrane helices (TM1-7), α5 helix of miniGq and the ligand of TUG-891 bound FFA4-miniGq complex are shown. The EM density is shown at 0.147 threshold.

**Figure S3.**
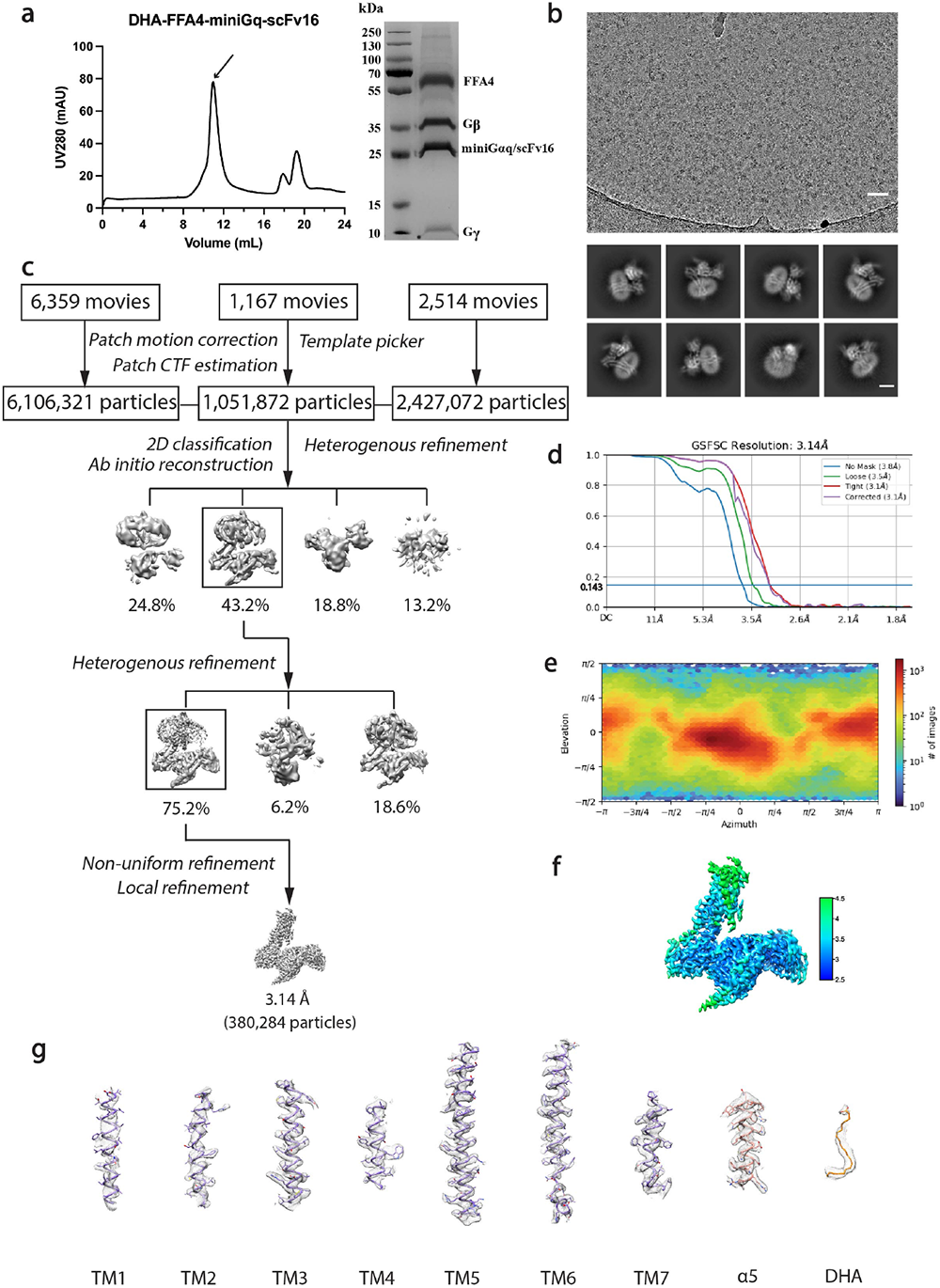
Purification of the FFA4-miniGq complex with DHA and cryo-EM data processing. **(a)** Size-exclusion chromatography profile and SDS-PAGE analysis of the purified DHA-FFA4-miniGq complex. **(b)** Representative cryo-EM micrograph (scale bar: 50 nm) and 2D class averages (scale bar: 5 nm). **(c)** Cryo-EM image processing workflow for the DHA-FFA4-miniGq complex. **(d)** Gold-standard Fourier shell correlation (FSC) curve showing an overall resolution is 3.14 Å at FSC=0.143. **(e)** Angular distribution of the particles used in the final reconstruction. **(f)** Density map according to local resolution estimation. **(g)** Cryo-EM density maps and models of the seven transmembrane helices (TM1-7), α5 helix of miniGq and the ligand of DHA bound FFA4-miniGq complex are shown. The EM density is shown at 0.132 threshold.

**Figure S4.**
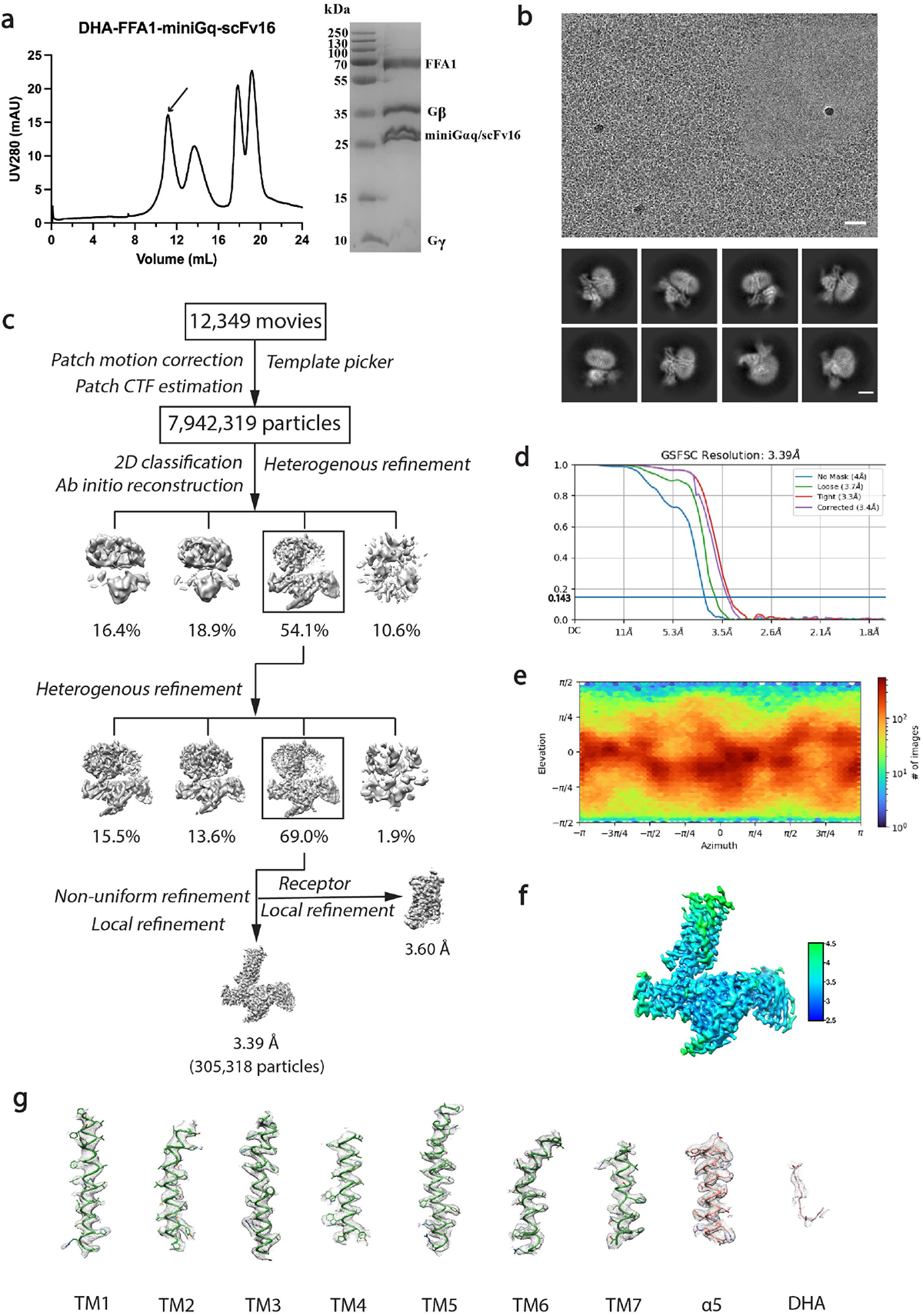
Purification of the FFA1-miniGq complex with DHA and cryo-EM data processing. **(a)** Size-exclusion chromatography profile and SDS-PAGE analysis of the purified DHA-FFA1-miniGq complex. **(b)** Representative cryo-EM micrograph (scale bar: 50 nm) and 2D class averages (scale bar: 5 nm). **(c)** Cryo-EM image processing workflow for the DHA-FFA1-miniGq complex. **(d)** Gold-standard Fourier shell correlation (FSC) curve showing an overall resolution is 3.39 Å at FSC=0.143. **(e)** Angular distribution of the particles used in the final reconstruction. **(f)** Density map according to local resolution estimation. **(g)** Cryo-EM density maps and models of the seven transmembrane helices (TM1-7), α5 helix of miniGq and the ligand of DHA bound FFA1-miniGq complex are shown. The EM density is shown at 0.128 threshold.

**Figure S5.**
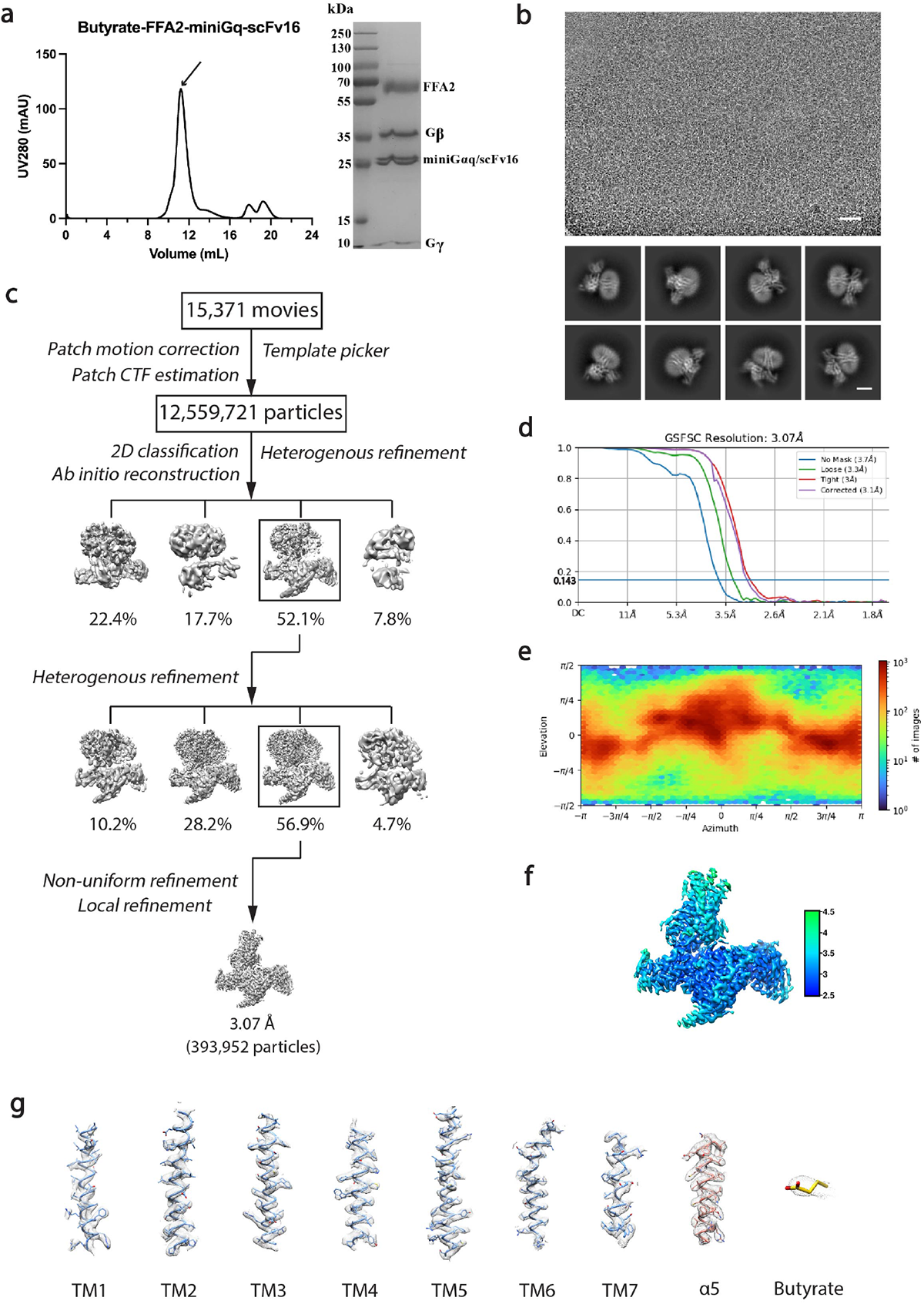
Purification of the FFA2-miniGq complex with butyrate and cryo-EM data processing. **(a)** Size-exclusion chromatography profile and SDS-PAGE analysis of the purified Butyrate-FFA2-miniGq complex. **(b)** Representative cryo-EM micrograph (scale bar: 50 nm) and 2D class averages (scale bar: 5 nm). **(c)** Cryo-EM image processing workflow for the Butyrate-FFA2-miniGq complex. **(d)** Gold-standard Fourier shell correlation (FSC) curve showing an overall resolution is 3.07 Å at FSC=0.143. **(e)** Angular distribution of the particles used in the final reconstruction. **(f)** Density map according to local resolution estimation. **(g)** Cryo-EM density maps and models of the seven transmembrane helices (TM1-7), α5 helix of miniGq and the ligand of Butyrate bound FFA2-miniGq complex are shown. The EM density is shown at 0.152 threshold.

**Figure S6.**
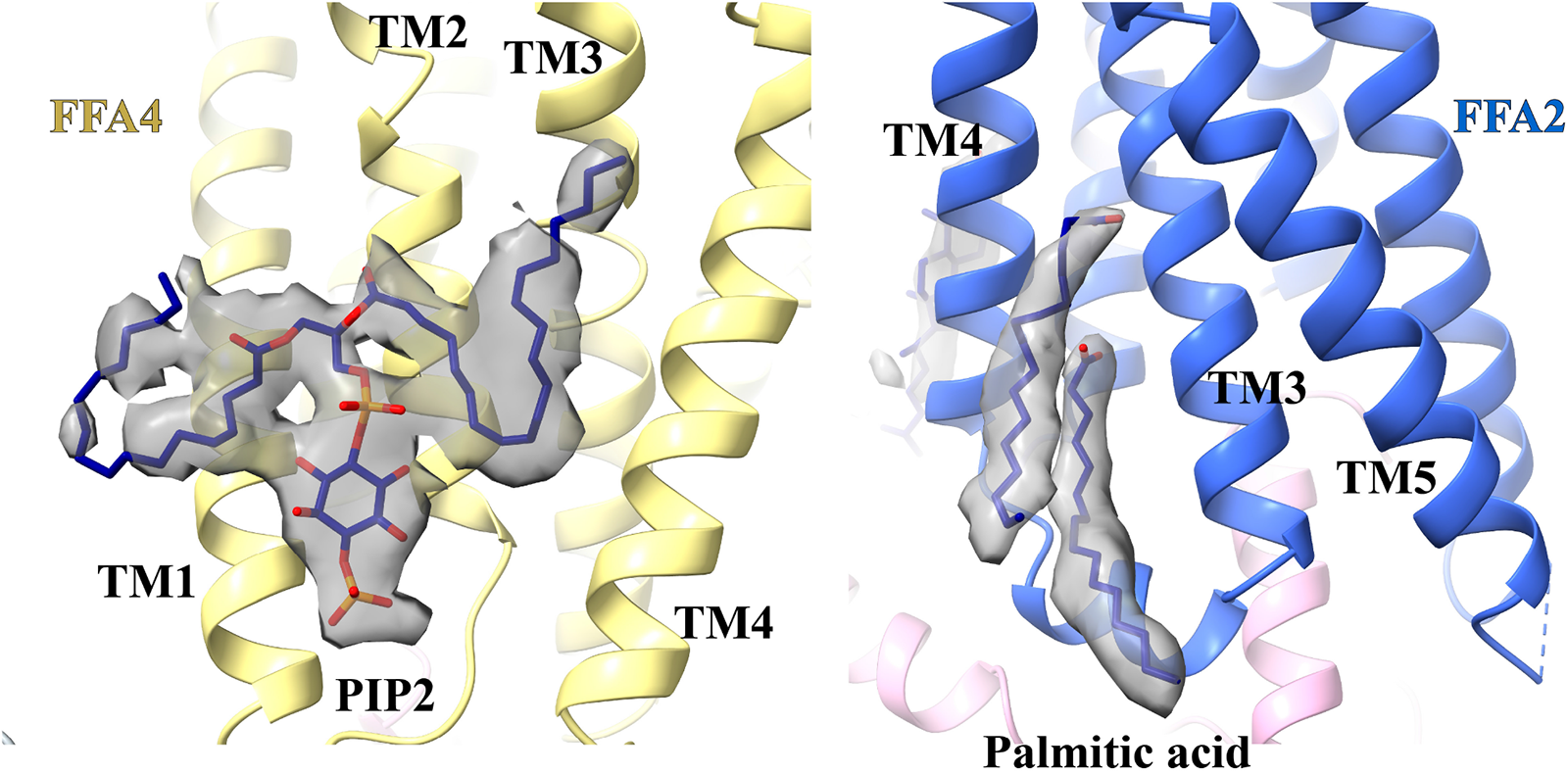
Lipid molecules surrounding 7TMs of FFA2 (blue) and FFA4 (yellow). PIP2 and palmitic acid molecules shown as dark blue sticks are modeled to fit the cryo-EM density (dark grey). The cryo-EM density of PIP2 is contoured at level 0.12. The cryo-EM density of palmitic acid is contoured at level 0.15.

**Figure S7.**
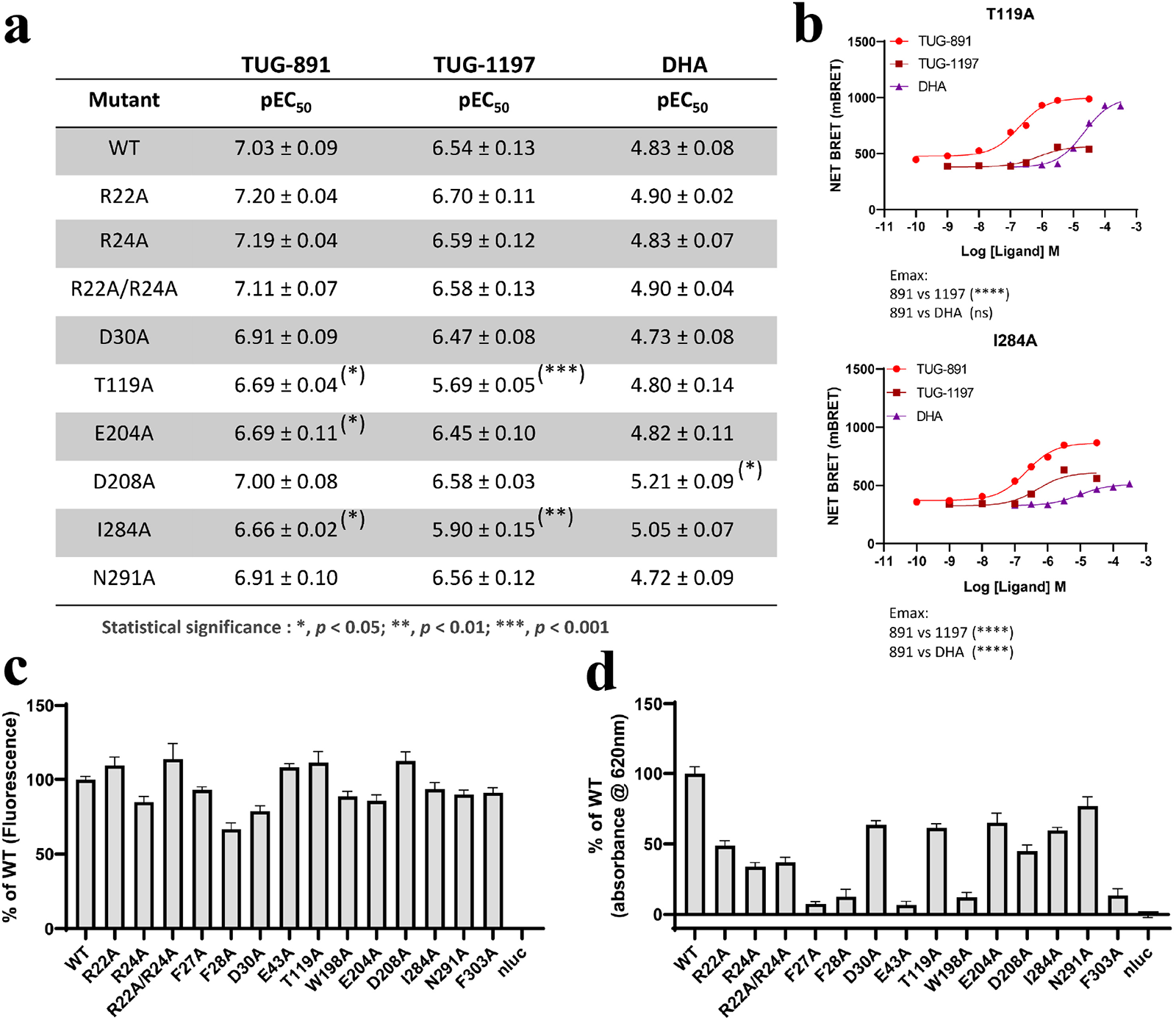
Mutagenesis studies on FFA4. **(a)** Point mutants of FFA4 were compared to wild type (WT) in arrestin 3 interaction studies to explore potential alterations in potency for TUG-891, TUG-1197 and DHA. Data are means +/-S.E.M. for n = 3 or more. Significantly different from wild type at p < 0.05 *, < 0.01** and 0.001 ***. **(b)** Concentration-response curves for TUG-891, TUG-1197 and DHA at T119A (**upper panel**) and I284A (**lower panel**) highlights the reduced efficacy of DHA and TUG1197 at I284A but only for TUG-1197 at T119A. **(c)** whilst all FFA4-eYFP mutants were expressed effectively as measured by eYFP fluorescence, **(d)** only a subset were effectively delivered to the cell surface as quantified by ELISA measurements. We were thus unable to define the effect of mutation to Ala of each of F27, F28 and E43 (see text for details).

**Figure S8.**
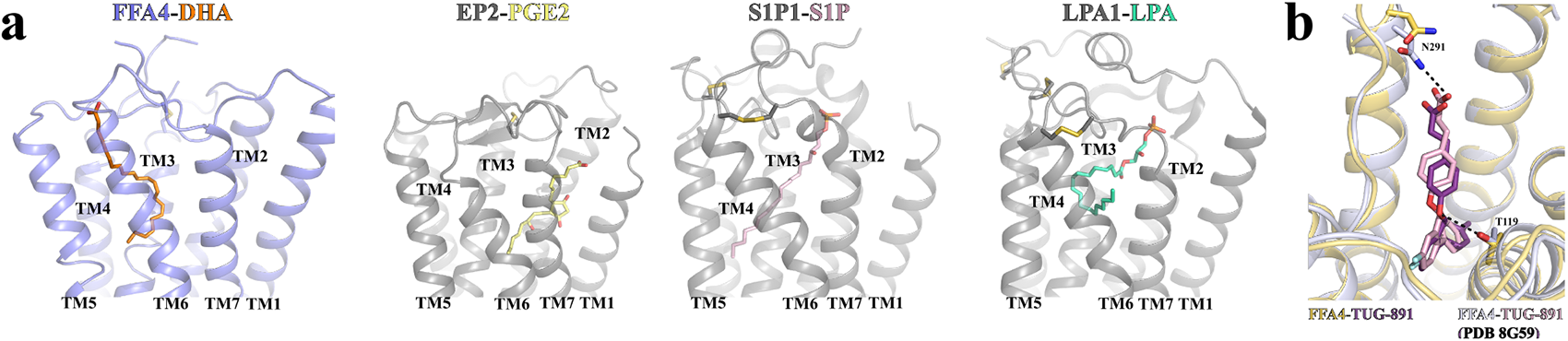
Structural comparison of FFA4 with (a) other lipid GPCRs and (b) published structure of FFA4 with TUG-891. EP2 is the receptor for the prostaglandin E2 (PGE2). S1P1 and LPA1 are receptors for the lysophospholipids S1P and LPA, respectively. The PDB IDs of the structures of EP2-PGE2, S1P1-S1P, and LPA1-LPA are 7XC2, 7TD3, and 7TD0, respectively. TUG-891 in our structure and in the published structure (PDB ID 8G59) is colored purple and pink, respectively.

**Figure S9.**
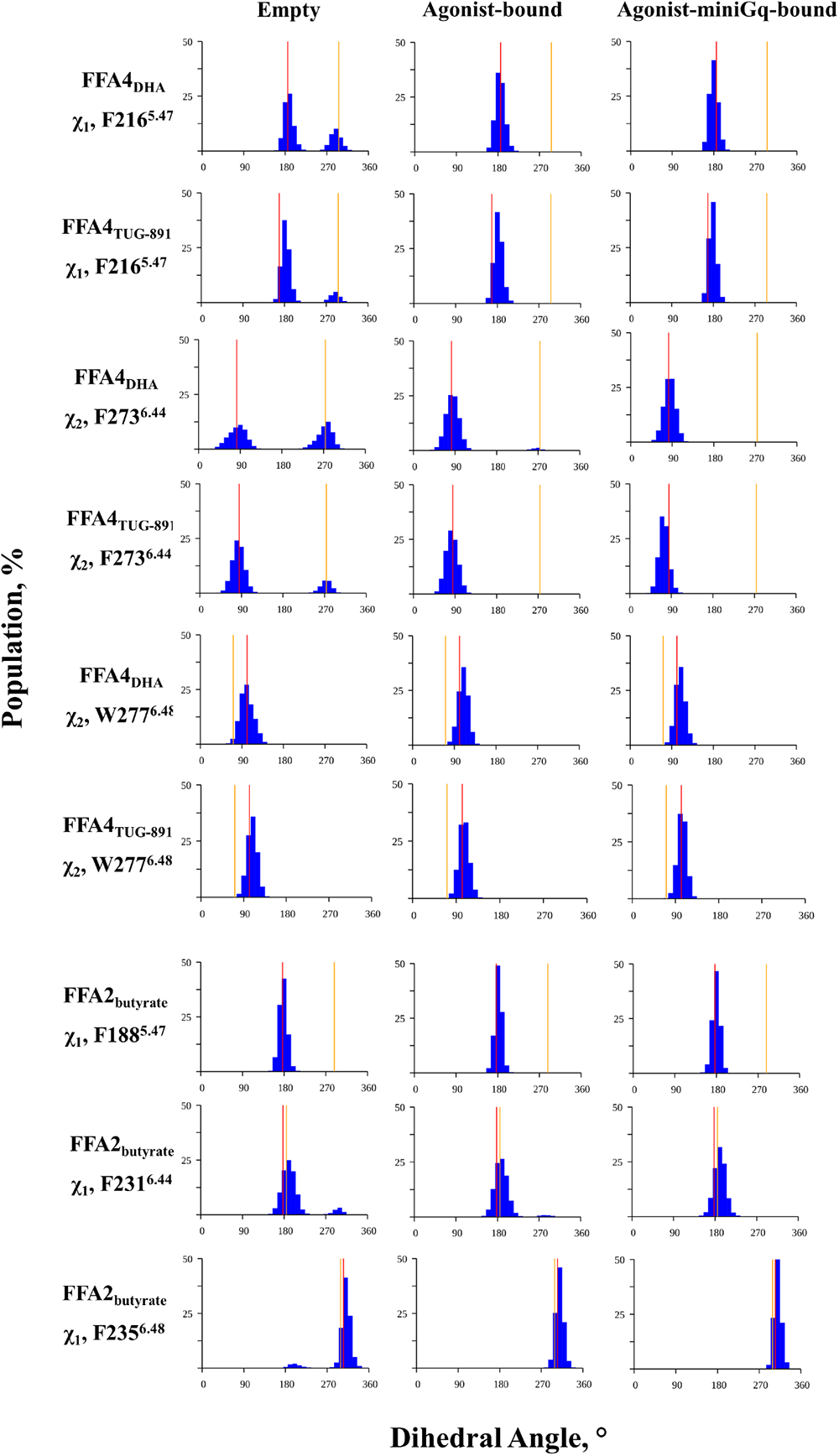
Dihedral angle population in MD simulations. The dihedral χ_1_ or χ_2_ angles of the aromatic residues in positions 5.47, 6.44, 6.45 and 6.48 are shown from the three 1 μs MD simulations of FFA4 and FFA2 in the empty, agonist- and agonist-miniG_q_-bound forms. The populations are calculated with bin width of 10°. The red and orange lines show a χ_1_ / χ_2_ value observed in the cryoEM active and Alphafold inactive structures, respectively.

**Figure S10.**
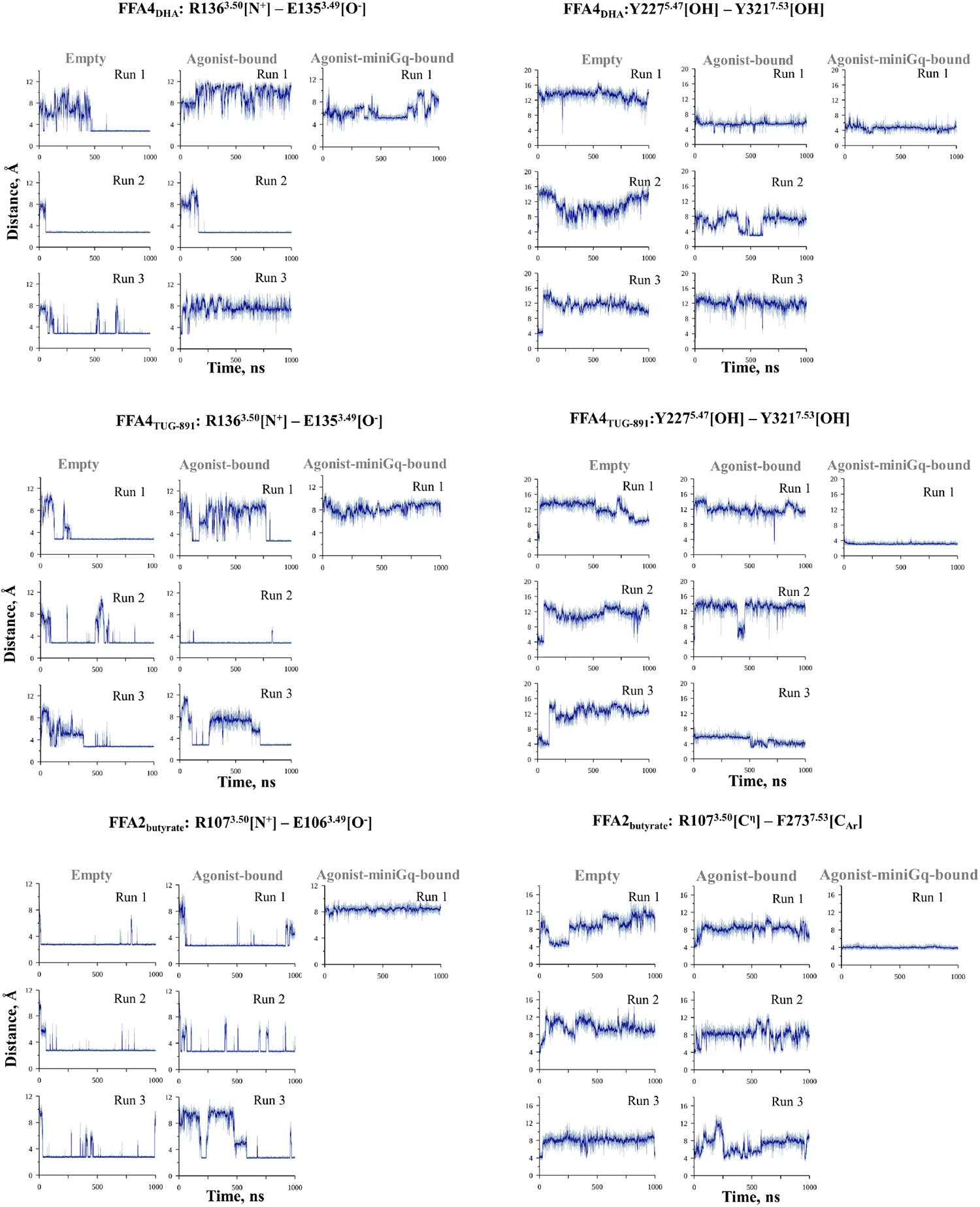
Evolution of atom distances in MD simulations. The distance between atoms of conserved amino acids known to be important for GPCR activation are shown from the three 1 μs MD simulations of FFA4 and FFA2 in the empty, agonist- and agonist-miniG_q_-bound forms. The sets of atoms used to measure the minimum inter-atomic distance in each frame are given in brackets. The thin line represents the raw values while bold line represents the shifting average of 10 consecutive frames.

**Figure S11.**
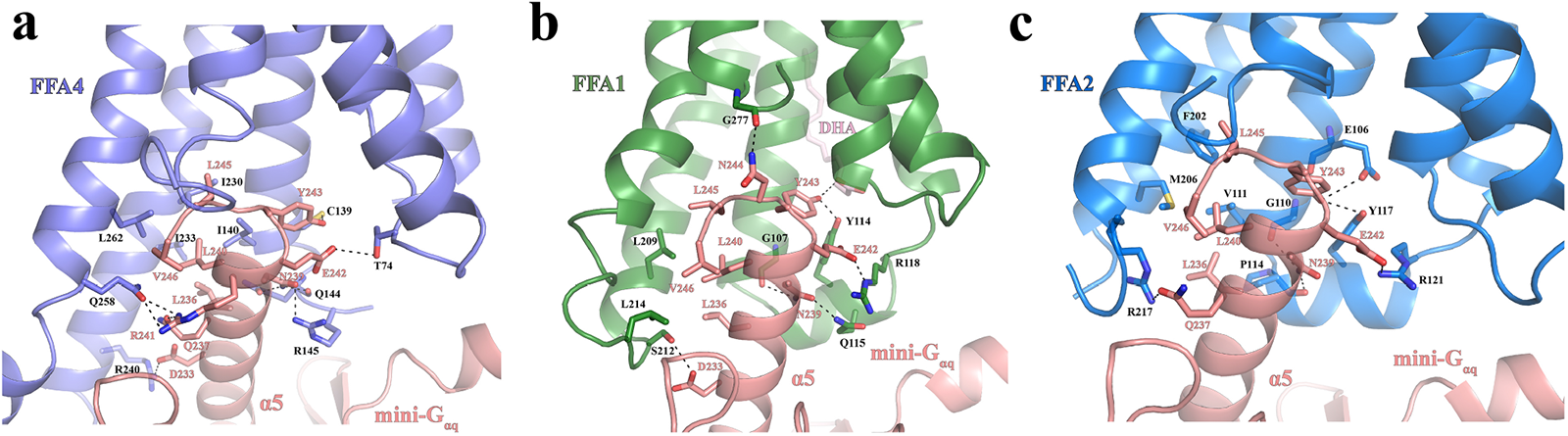
Interactions between FFAs and the α5 helix of mini-G_αq_. **(a)** Interactions between FFA4 and the α5 helix of mini-G_αq_. Specifically, hydrophobic interactions form among the mini-G_αq_ residues L236, L240, L245, and V246 and the FFA4 residues I140^3.54^, I230^5.61^, I233^5.64^, and L262^6.33^. In addition, Y243, E242, and D233 of mini-G_αq_ form polar interactions with the side chains of FFA4 residues C139^3.53^, T74^2.39^, and R240^5.71^, respectively. Furthermore, N239 of mini-G_αq_ forms polar interactions with FFA4 residues R145^ICL2^ and Q144 ^ICL2^, whereas R241 and Q237 of mini-G_αq_ form a polar interaction network with FFA4 residue Q258 ^6.29^. **(b)** Interactions between FFA1 and the α5 helix of mini-G_αq_. Specifically, hydrophobic interactions form among the mini-G_αq_ residues L236, L240, L245, and V246 and the FFA1 residues L209^5.65^ and L214^ICL3^. Y243, E242, N239, and D233 of mini-G_αq_ form polar interactions with the side chains of FFA1 residues Y114^ICL2^, R118^ICL2^, Q115^ICL2^, and S212^ICL3^, respectively. Furthermore, N244 and N239 of mini-G_αq_ forms polar interactions with the carbonyl groups of FFA1 residues G277^7.54^ and G107^3.53^, respectively. Y243 of mini-G_αq_ also forms a hydrogen bond with DHA that may be bound at ‘Site 2’. **(c)** Interactions between FFA2 and the α5 helix of mini-G_αq_. Specifically, hydrophobic interactions form among the mini-G_αq_ residues L236, L240, L245, and V246 and the FFA2 residues M206^5.65^ and F202^5.61^. In addition, E242 and Q237 of mini-G_αq_ form polar interactions with the side chains of FFA2 residues R121^ICL2^ and R217^6.30^, respectively. Furthermore, Y243 of mini-G_αq_ forms polar interactions with FFA2 residues E106^3.49^ and Y117^ICL2^, whereas 239 of mini-G_αq_ form polar interactions with the mainchain carbonyl groups of FFA4 residue G110^3.53^ and P114_ICL2._

**Figure S12.**
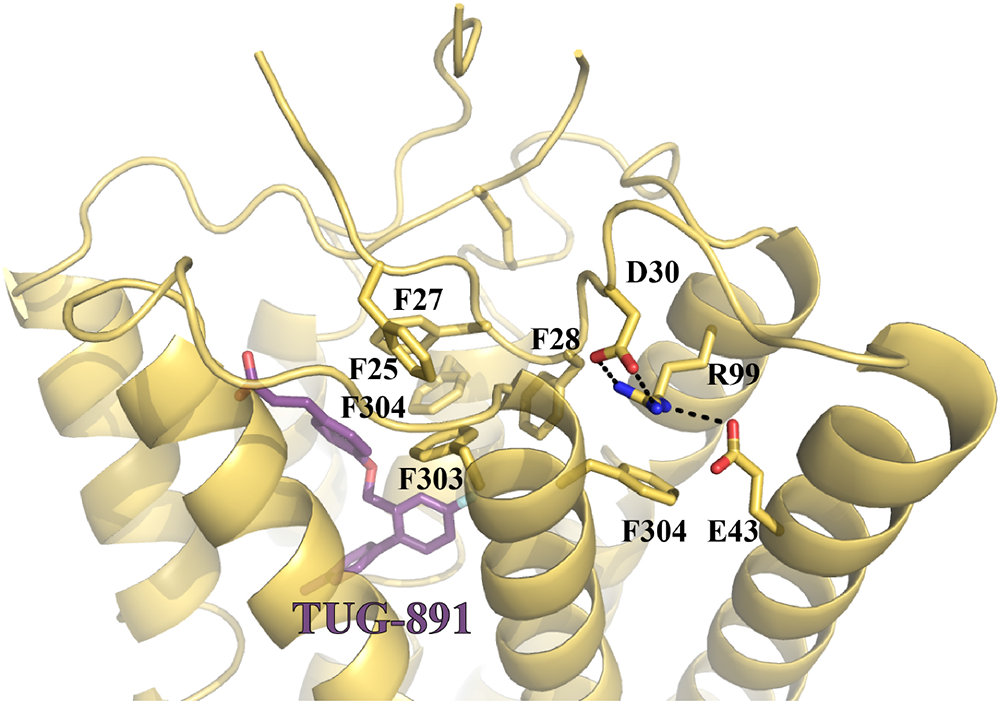
Interactions mediated by R99 of FFA4 and nearby aromatic network. In contrast to initial predictions based on homology modeling and mutagenesis R99^2.64^ does not interact directly with the carboxylate ofTUG-891. Rather it acts to shape the binding pocket through interactions with D30^N^ and E43^1.35^ and a cation-π interaction with F304^7.36^.

**Table S1.** Cryo-EM data collection and refinement statistics 1.

|  | (EMD-41007, PDB 8T3O) | (EMD-41008, PDB 8T3Q) |
| --- | --- | --- |
| Data collection and processing | TUG-891-FFA4-miniGq | DHA-FFA4-miniGq |
| Magnification | 105,000 | 105,000 |
| Voltage (kV) | 300 | 300 |
| Electron exposure (e<sup>-</sup>/Å<sup>2</sup>) | 55 | 55 |
| Defocus range (μm) | -1.0 to -1.8 | -1.0 to -1.8 |
| Pixel size (Å) | 0.828 | 0.828 |
| Symmetry imposed | C1 | C1 |
| Initial particle images (no.) | 4,928,436 | 9,585,265 |
| Final particle images (no.) | 391,203 | 380,284 |
| Map resolution (Å) | 3.06 | 3.14 |
| FSC threshold | 0.143 | 0.143 |
| Map resolution range (Å) | 2.5-4.5 | 2.5-4.5 |
| Refinement |  |  |
| Model resolution (Å) | 3.0 | 3.1 |
| FSC threshold | 0.143 | 0.143 |
| Model composition |  |  |
| Non-hydrogen atoms | 9112 | 9032 |
| Protein residues | 1158 | 1158 |
| Ligand | 1 | 1 |
| R.m.s. deviations |  |  |
| Bond lengths (Å) | 0.005 | 0.008 |
| Bond angles (°) | 0.802 | 1.160 |
| Validation |  |  |
| MolProbity score | 1.65 | 2.00 |
| Clashscore | 8.65 | 11.84 |
| Rotamer outliers (%) | 0.21 | 1.44 |
| Ramachandran plot |  |  |
| Favored (%) | 96.93 | 95.79 |
| Allowed (%) | 3.07 | 4.21 |
| Disallowed (%) | 0 | 0 |

**Table S2.** Cryo-EM data collection and refinement statistics 2.

|  | (EMD-41013, PDB 8T3V)/(EMD-41014) | (EMD-41010, PDB 8T3S) |
| --- | --- | --- |
| Data collection and processing | DHA-FFA1-miniGq/Local | butyrate-FFA2-miniGq |
| Magnification | 105,000 | 105,000 |
| Voltage (kV) | 300 | 300 |
| Electron exposure (e<sup>-</sup>/Å<sup>2</sup>) | 61.6 | 61.6 |
| Defocus range (μm) | -0.8 to -2.5 | -0.8 to -2.5 |
| Pixel size (Å) | 0.826 | 0.826 |
| Symmetry imposed | C1 | C1 |
| Initial particle images (no.) | 7,942,319 | 12,559,721 |
| Final particle images (no.) | 305,318 | 393,952 |
| Map resolution (Å) | 3.39/3.6 | 3.07 |
| FSC threshold | 0.143 | 0.143 |
| Map resolution range (Å) | 2.5-4.5 | 2.5-4.5 |
| Refinement |  |  |
| Model resolution (Å) | 3.4 | 3.0 |
| FSC threshold | 0.143 | 0.143 |
| Model composition |  |  |
| Non-hydrogen atoms | 8745 | 8772 |
| Protein residues | 1136 | 1119 |
| Ligand | 1 | 1 |
| R.m.s. deviations |  |  |
| Bond lengths (Å) | 0.007 | 0.007 |
| Bond angles (°) | 0.873 | 0.797 |
| Validation |  |  |
| MolProbity score | 1.69 | 1.60 |
| Clashscore | 11.48 | 9.17 |
| Rotamer outliers (%) | 0.44 | 0.53 |
| Ramachandran plot |  |  |
| Favored (%) | 97.41 | 97.45 |
| Allowed (%) | 2.59 | 2.55 |
| Disallowed (%) | 0 | 0 |

**Table S3.** The average root mean square deviation and fluctuation (RMSD and RMSF) of the receptor systems. RMSD and RMSF values were calculated for the receptor Cα atoms, the 7-transmembrane bundle (7TMB) Cα atoms and the ligand non-hydrogen atoms (Ligand) from the three 1µ MD simulations for receptor-only systems and one 1 μs MD simulation for receptor-miniGq systems. RMSD values were averaged within each replica and combined between different replicas to yield the mean ± standard deviation values given below. RMSF values are also given as mean ± standard deviation between all replicas.

| Receptor Systems | RMSD, Å |  |  | RMSF, Å |  |  |
| --- | --- | --- | --- | --- | --- | --- |
| | Cα | 7TMB-Cα | Ligand | Cα | 7TMB-Cα | Ligand |
| FFA4<sub>DHA</sub> | 3.6 ± 0.2 | 1.3 ± 0.0 | 3.9 ± 1.1 | 1.4 ± 0.1 | 0.7 ± 0.0 | 1.9 ± 0.2 |
| FFA4<sub>empty</sub> | 3.7 ± 0.1 | 1.5 ± 0.0 | | 1.6 ± 0.2 | 0.8 ± 0.1 | |
| FFA4<sub>DHA_Gq</sub> | 3.4 | 1.2 | 4.0 | 1.4 | 0.7 | 2.2 |
| FFA4<sub>TUG-891</sub> | 3.7 ± 0.3 | 1.2 ± 0.0 | 1.9 ± 0.2 | 1.3 ± 0.1 | 0.7 ± 0.0 | 1.0 ± 0.1 |
| FFA4<sub>empty_TUG-891</sub> | 3.2 ± 0.4 | 1.3 ± 0.1 | | 1.5 ± 0.2 | 0.8 ± 0.1 | |
| FFA4<sub>TUG-891_Gq</sub> | 2.8± | 1.2 | 1.9 | 1.1 | 0.6 | 1.0 |
| FFA2<sub>butyrate</sub> | 2.6 ± 0.2 | 1.1 ± 0.0 | 2.2 ± 0.4 | 1.2 ± 0.0 | 0.7 ± 0.0 | 1.6 ± 0.3 |
| FFA2<sub>empty</sub> | 2.8 ± 0.1 | 1.3 ± 0.2 | | 1.2 ± 0.1 | 0.8 ± 0.1 | |
| FFA2<sub>butyrate_Gq</sub> | 2.2 | 1.2 | 1.9 | 0.9 | 0.6 | 1.5 |

**Table S4.**
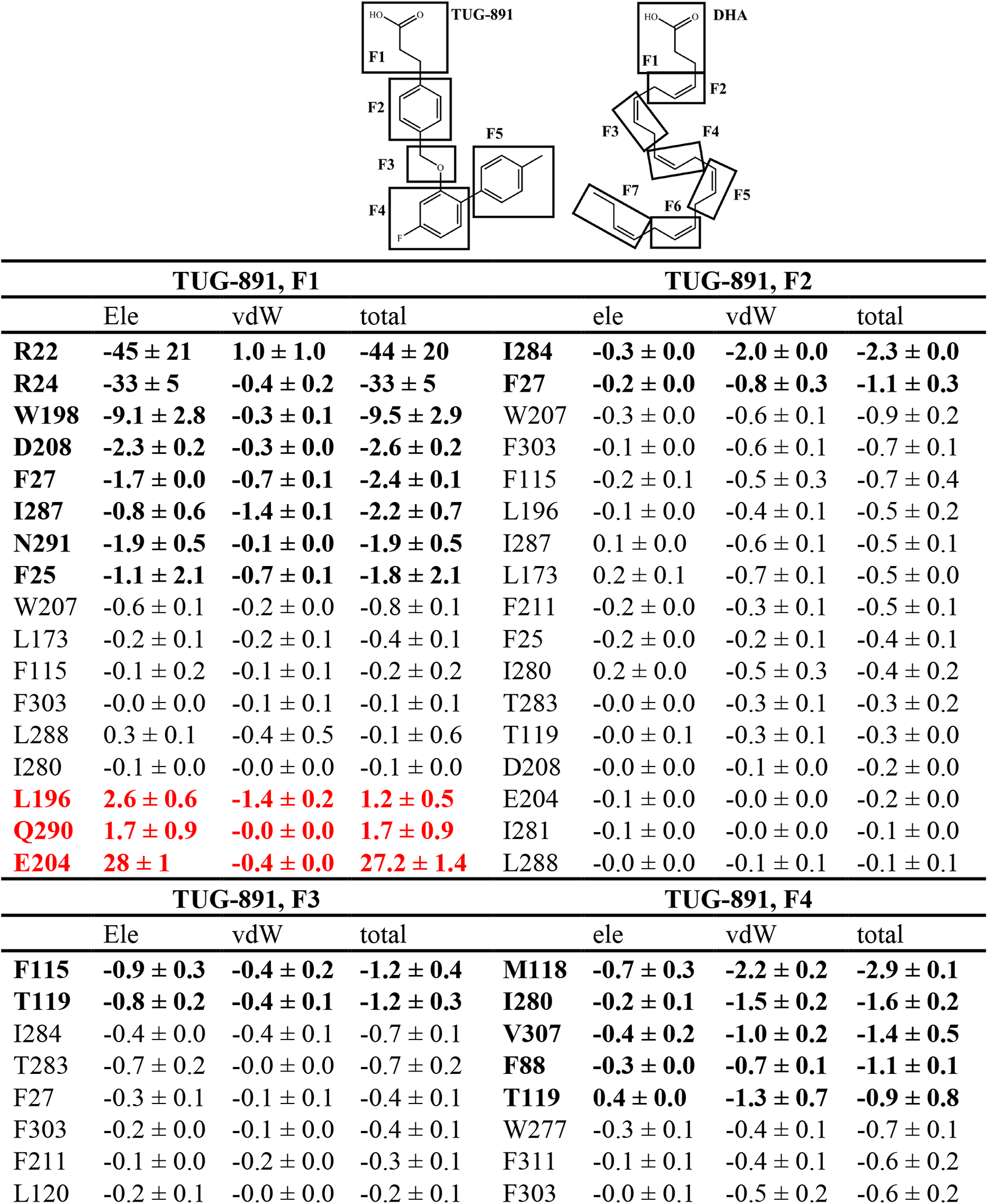
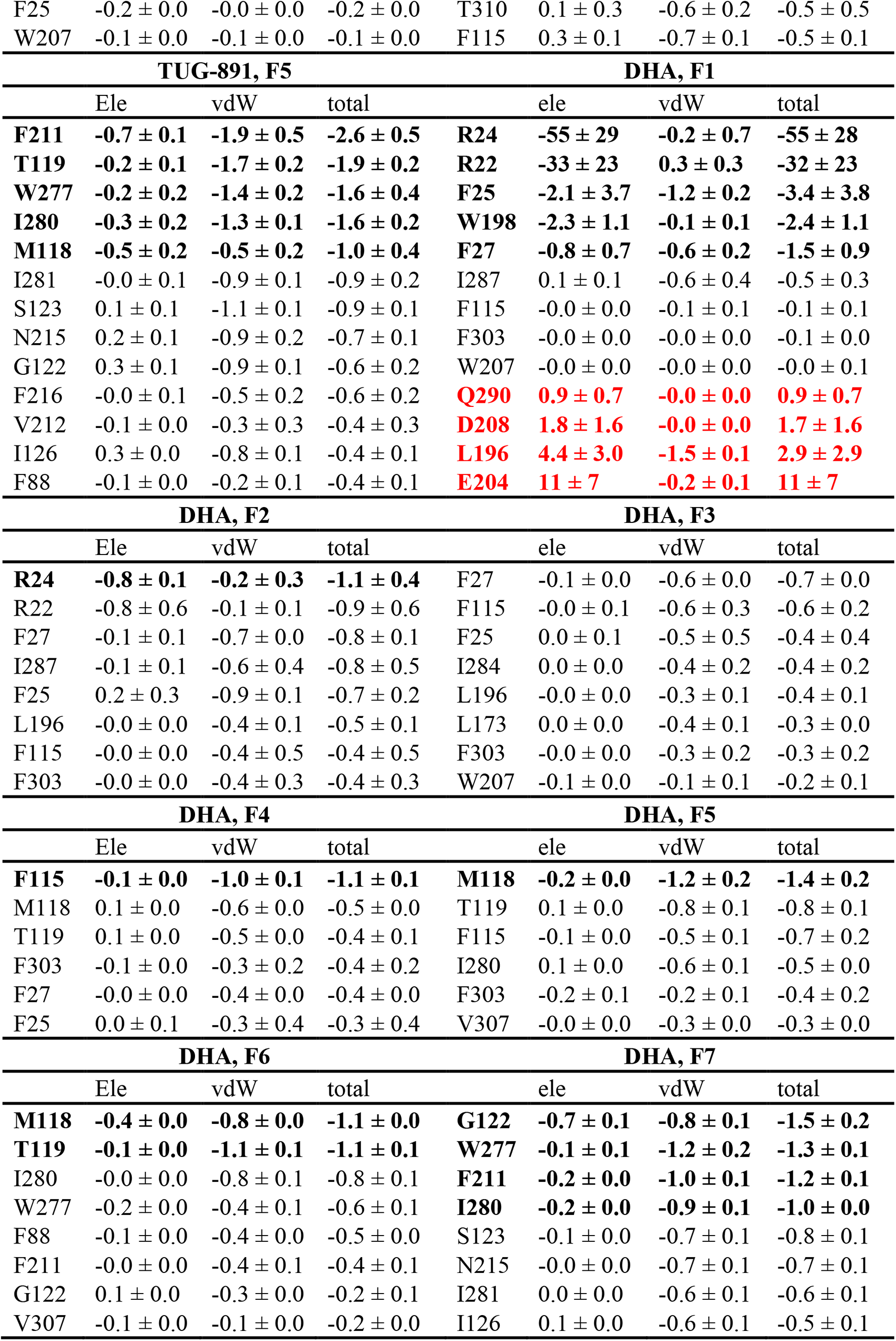
The average ligand fragment-residue interaction energy. The interaction energy involving electrostatic (ele) and van der Waals (vdW) components in kcal/mol was calculated from three 1µ MD simulations of the FFA4-agonist complexes. The fragmentation of the agonists is shown below. The residues having noticeable interactions with a given ligand fragment, i.e. having total energy below -1 kcal/mol or above 1 kcal/mol are shown in bold. Among them, residues having repulsive interactions, i.e. positive total energy, are colored red. The values were averaged within each replica and combined between different replicas to yield the mean ± standard deviation values given below.

